# Neural manifold under plasticity in a goal driven learning behaviour

**DOI:** 10.1101/2020.02.21.959163

**Authors:** Barbara Feulner, Claudia Clopath

**Affiliations:** Imperial College London

## Abstract

Neural activity is often low dimensional and dominated by only a few prominent neural covariation patterns. It has been hypothesised that these covariation patterns could form the building blocks used for fast and flexible motor control. Supporting this idea, recent experiments have shown that monkeys can learn to adapt their neural activity in motor cortex on a timescale of minutes, given that the change lies within the original low-dimensional subspace, also called neural manifold. However, the neural mechanism underlying this within-manifold adaptation remains unknown. Here, we show in a computational model that modification of recurrent weights, driven by a learned feedback signal, can account for the observed behavioural difference between within- and outside-manifold learning. Our findings give a new perspective, showing that recurrent weight changes do not necessarily lead to change in the neural manifold. On the contrary, successful learning is naturally constrained to a common subspace.

## Introduction

The dynamics of single neurons within a given brain circuit are not independent, but highly correlated. Moreover, neural activity is often dominated by only a very low number of distinct correlation patterns [Churchland et al., 2012, Mante et al., 2013, Kaufman et al., 2014, Cunningham and Yu, 2014, Gao and Ganguli, 2015, Williamson et al., 2016, Pang et al., 2016, Mazzucato et al., 2016, Michaels et al., 2016, Elsayed and Cunningham, 2017, Gallego et al., 2017, Williams et al., 2018, Rus, 2018, Perich et al., 2018, Semedo et al., 2019]. This implies that there exists a low-dimensional manifold in the high-dimensional population activity space, which most of the variance of the neural activity is confined to. Why such low-dimensional dynamics is observed in the brain remains unclear. One potential reason is that most experimental designs are inherently low-dimensional, and therefore bias the observed neural activity [Gao et al., 2017]. Alternatively, it has been shown that low-dimensional dynamics can arise from structured connectivity within the network [Sussillo and Abbott, 2009, Laje and Buonomano, 2013, Hu et al., 2014, Hennequin et al., 2014, Aljadeff et al., 2016, Rivkind and Barak, 2017, Mastrogiuseppe and Ostojic, 2018, DePasquale et al., 2018, Recanatesi et al., 2019].

The functional implications of low-dimensional neural dynamics are also unclear [Rigotti et al., 2013, Rule et al., 2019]. For example, preparatory and movement activity lie in orthogonal subspaces, which can serve as a gating mechanism for movement initiation [Kaufman et al., 2014, Elsayed et al., 2016]. Furthermore, the same manifold might underlie multiple skilled wrist and reach-to-grasp movements [Gallego et al., 2018], as well as the same task over a long timescale [Gallego et al., 2020]. These results suggest that there is one stable manifold for motor control, and that learning and performing different, but similar, movements are happening within this scaffold [Gallego et al., 2017].

Using a brain-computer interface (BCI) [Golub et al., 2016] in monkeys, Sadtler et al. showed that it is possible to adapt neural activity in motor cortex, if the new activity pattern lies within the original manifold, but not if it lies outside of it [Sadtler et al., 2014]. They could test this by introducing two kinds of perturbations to a BCI, which was previously trained to decode two-dimensional cursor dynamics from neural activity. The monkeys had to adapt their neural activity in motor cortex to produce the correct cursor dynamics, given the perturbed BCI mapping. Interestingly, the timescale of within-manifold learning is in the order of minutes to hours [Sadtler et al., 2014]. In contrast, outside-manifold learning is not possible within a single session, but instead requires progressive training over multiple days [Oby et al., 2019]. Due to the timescale difference, it has been hypothesized that within-manifold learning is related to fast motor adaption and outside-manifold learning is related to skill learning. Yet, the underlying neural mechanisms for both types of learning remain unknown.

We used computational modelling to study the relationship between neural manifold and learning. As motor learning can drive network rewiring in motor cortex on a short timescale [Xu et al., 2009, Biane et al., 2019, Tavor et al., 2019, Ohbayashi, 2020], we wanted to test whether local network rewiring can account for within-manifold, as well as outside-manifold learning. To that end, we implemented an in-silico version of the BCI experiment in Sadtler et al., where motor cortex activity is simulated by a recurrent neural network (RNN) [Sussillo and Abbott, 2009, Laje and Buonomano, 2013, DePasquale et al., 2018]. We showed that an ideal observer training algorithm is able to learn within- and outside-manifold perturbations equally well. One component of the algorithm is that errors in the produced cursor dynamics are translated to errors in single neuron activity. This form of feedback signal is used to adapt the local connections within the network in order to minimize the observed error. As the correct feedback signal is not a priori given in the biological system, we subsequently tested more realistic feedback scenarios. By disrupting the correct feedback signal, we found that within-manifold learning is more robust to erroneous or sparse feedback signals. Furthermore, learning the feedback signal from scratch was only possible for within-manifold perturbations. To enable feedback learning for outside-manifold perturbations we had to introduce an incremental strategy, which resembles the progressive training used in experiments [Oby et al., 2019].

## Results

### Recurrent neural network (RNN) performing a center-out-reach task

To investigate the potential difference between within- and outside-manifold learning, we implemented an in-silico version of a brain-computer interface (BCI) experiment previously done with monkeys [Sadtler et al., 2014]. Instead of measuring neural activity in monkey motor cortex, we implemented a recurrent neural network (RNN) which was trained to do a similar center-out-reach task as the monkey had to perform (Fig.1A). After the initial training phase, we perturbed the BCI mapping to either trigger a within- or outside-manifold change (Fig.1C & D). We then retrained the recurrent network. This pipeline allowed us to test to what extent reorganization in the local network can explain the behaviourally observed differences in within- and outside-manifold learning.

**Figure 1:**
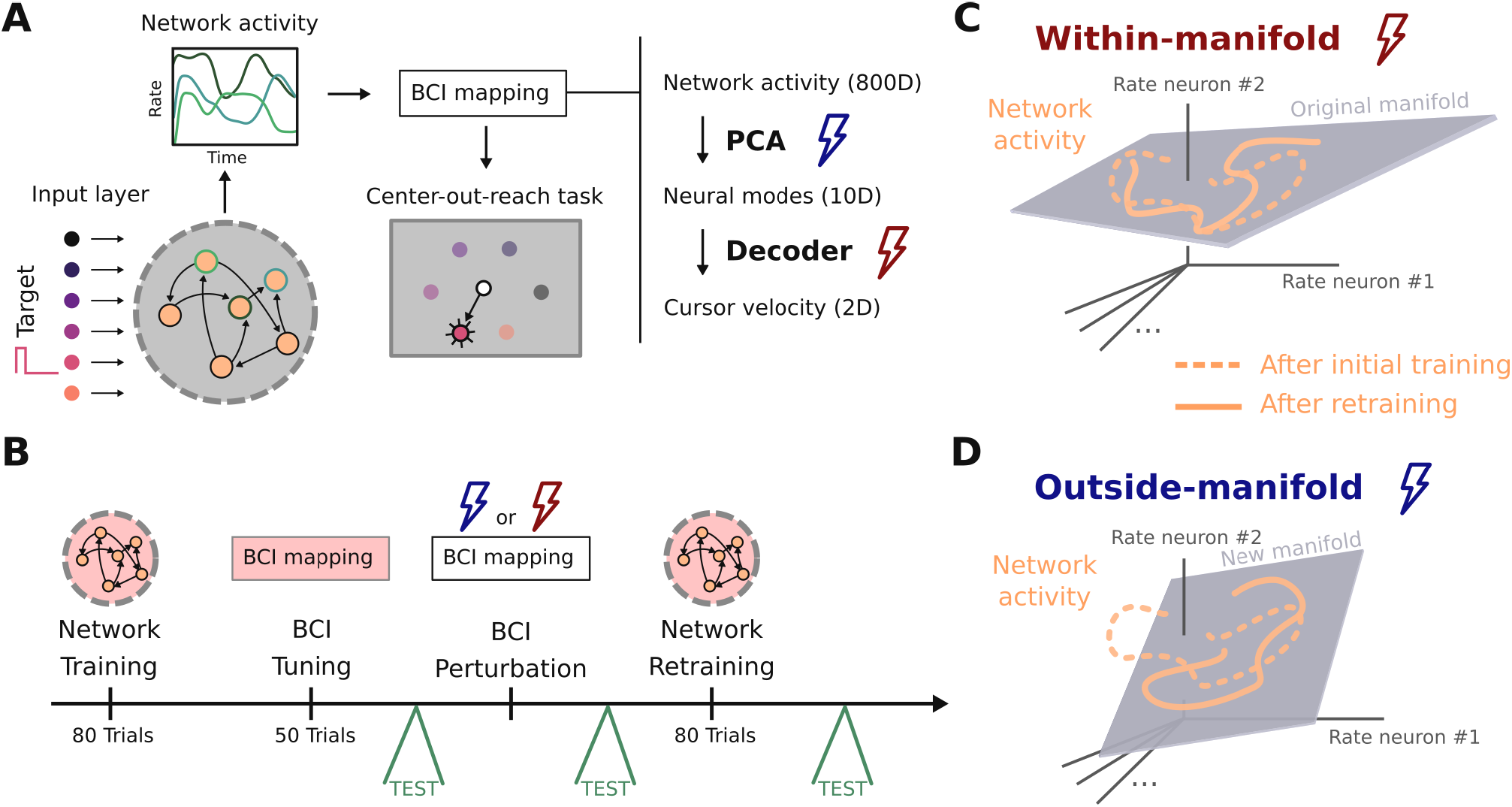
Recurrent neural network (RNN) performing a center-out-reach task. (A) In order to study learning within and outside of the neural manifold, we used a brain-computer interface (BCI) setup similar to [Sadtler et al., 2014]. A recurrent neural network serves as generator of network activity which is then used via a BCI setup to control a cursor in a two-dimensional center-out-reach task. The key step in our simulation is to perturb the initial BCI mapping and let the recurrent network learn to compensate for that. (B) The simulation design. (C and D) In order to compensate the BCI perturbation the network needs to produce new dynamics. For a within-manifold perturbation (C) the new target activity is constrained to the same manifold as the original activity. For an outside-manifold perturbation (D) this is not the case, but the network activity has to explore new dimensions in state space.

### RNN with correct feedback signal can learn within- and outside-manifold equally well

In order to understand constraints on within- and outside-manifold learning, we firstly simulated our network with an ideal-observer feedback signal. This means that the task error, which is defined as the difference between target and produced cursor velocity, is correctly assigned to errors on the activity of single neurons in the network. The correct credit assignment can be achieved by a matrix multiplication with the pseudo-inverse of the BCI readout. By doing this, we intentionally ignored the fact that a biological system would need to learn this credit assignment first, before it can be used for retraining. Rather, we first focused on the restructuring of the recurrent connections. With this training method, our network reached high performance in the center-out-reach task after the initial training phase (Fig.2A1). To probe within- versus outside-manifold learning, we then applied the corresponding perturbation to the BCI mapping, which led to impaired cursor trajectories in both cases (Fig.2A2). Interestingly, using the same training method as during the initial training phase, we could retrain the network to adapt to the changed BCI mapping for within-manifold perturbation, as well as for outside-manifold perturbation. The cursor trajectories after the initial training phase and after the retraining phase looked similar for both types of perturbations (Fig.2A3). We quantified the performance by measuring the mean squared error between target and produced cursor velocities (Fig.2B). The average retraining performance confirms that within- and outside-manifold perturbations can equally well be counteracted and that this result is not constrained to a specific network initialization (Fig.2B), task setup (Fig.S1 & S5), or learning algorithm (Fig.S3 & S6). Together, these results indicate that there is no fundamental difference in learning new dynamics either within the original manifold or outside of it, as long as we focus solely on the learning of the recurrent weights.

**Figure 2:**
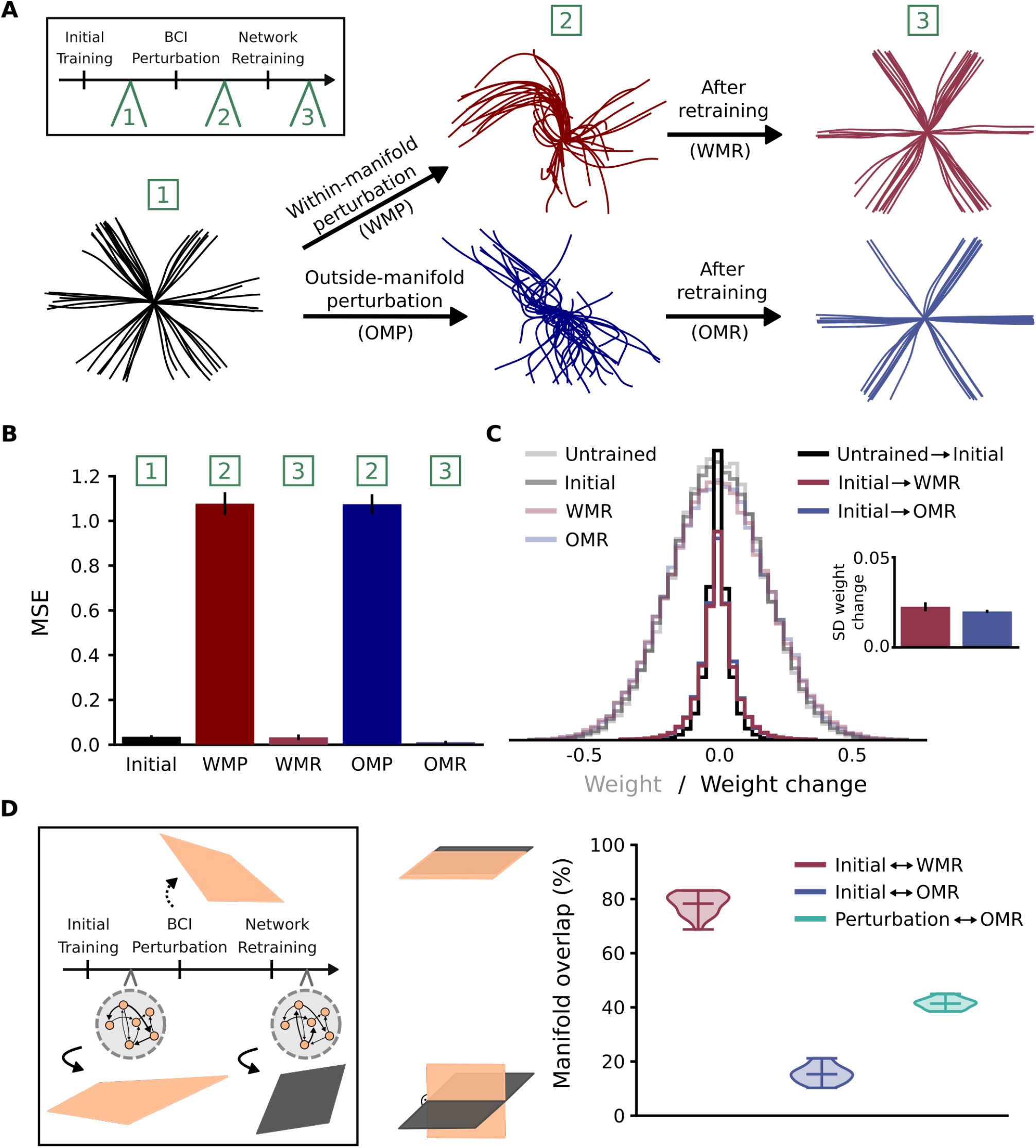
RNN with correct feedback signal can learn within- and outside-manifold equally well. (A) Cursor trajectories after initial training phase (on the left), after within- (WMP) or outside-manifold (OMP) perturbation (in the middle) and after within (WMR) or outside (OMR) retraining phase (on the right) for one example simulation. (B) Performance measured as mean squared error (MSE) between target cursor velocity and produced cursor velocity. Shown are the average simulation results for twenty randomly initialized networks, bars indicate the standard deviation across networks. (C) Weight distribution and weight change distribution during the experiment. The inlet shows the average standard deviation of weight change, bars indicate standard deviation across networks. (D) Manifold overlap between before and after retraining phase (red and blue), or between target manifold for outside-manifold perturbation and internal manifold after retraining phase (green).

Given biological constraints on the amount of affordable synaptic restructuring, we wanted to test whether there is a difference in weight change needed to compensate either within- or outside-manifold perturbations. To investigate this, we started with comparing the network connectivity before and after the retraining phase (Fig.2C). We found that there is no difference in the amount of weight change produced by either within- or outside-manifold retraining. In fact, the weight change distribution for both types of retraining is similar to the distribution obtained by comparing the weights before and after the initial training phase. These results show that retraining within or outside of the original manifold does not necessarily lead to a different amount of weight change.

In order to understand how our network achieves high performance after the adaptation phase, we analysed the underlying dynamics before and after adaptation. To quantify the dynamical changes, we recalculated the internal manifold after the retraining phase and compared it to the original manifold (Fig.2D). For within-manifold relearning, we found that the internal manifolds before and after relearning have high overlap as expected (Fig.2D red). Interestingly, learning is not equally distributed across all ten modes in the manifold, but concentrated on the most prominent ones (Fig.S7). For outside-manifold, we computed two measures: 1) how much the internal manifold after retraining still overlaps with the original one (Fig.2D blue), and 2) how the internal manifold after retraining has aligned to the one defined by the BCI perturbation (Fig.2D green). We found that although the task performance is high after retraining the network following an outside-manifold perturbation, the network dynamics do not completely align with the new readout defined by the altered BCI mapping. More detailed analysis showed that outside-manifold learning focuses on the few most prominent modes, which have initially the highest decoder values (Fig.S8 & 9). This demonstrates that it is not necessary that network dynamics completely realign with the altered BCI mapping to perform well in the task. Instead it is enough to have a sufficient amount of overlap, or match the most prominent modes.

### RNN with corrupted feedback signal or constraint on weight change can learn better within- than outside-manifold

Our results so far do not show a difference between within- and outside-manifold learning, in constrast to experimental observations [Sadtler et al., 2014]. To further investigate where this behaviourally observed difference potentially comes from, we next dropped the assumption of an ideal-observer feedback signal, which is biologically highly implausible, and tested the effect of different corrupted feedback versions on recurrent retraining. We started by looking at the effect of noisy feedback signals (Fig.3B). We simulated our experiment with different strengths of noise corruption in the feedback signal and compared the task performance after retraining. Interestingly, we found a difference between within- and outside-manifold learning, as within-manifold learning shows for a certain intermediate noise level better retraining performance than outside-manifold learning (Fig.3B top). Next, we looked at the effect of sparse feedback signals (Fig.3C). In this scenario, not all neurons in the recurrent network receive a feedback signal and therefore participate in the recurrent restructuring. We again found that within-manifold learning is achieving better performance with the same portion of feedback signals available (Fig.3C top). Finally, we tested the effect of sparse plastic weights within the recurrent network (Fig.3D). Also there we found that within-manifold learning is preferential. All together, these results show that within-manifold learning outperforms outside-manifold learning in the presence of either erroneous feedback signals, sparse feedback signals, or if there is a constrained number of plastic connections within the network.

**Figure 3:**
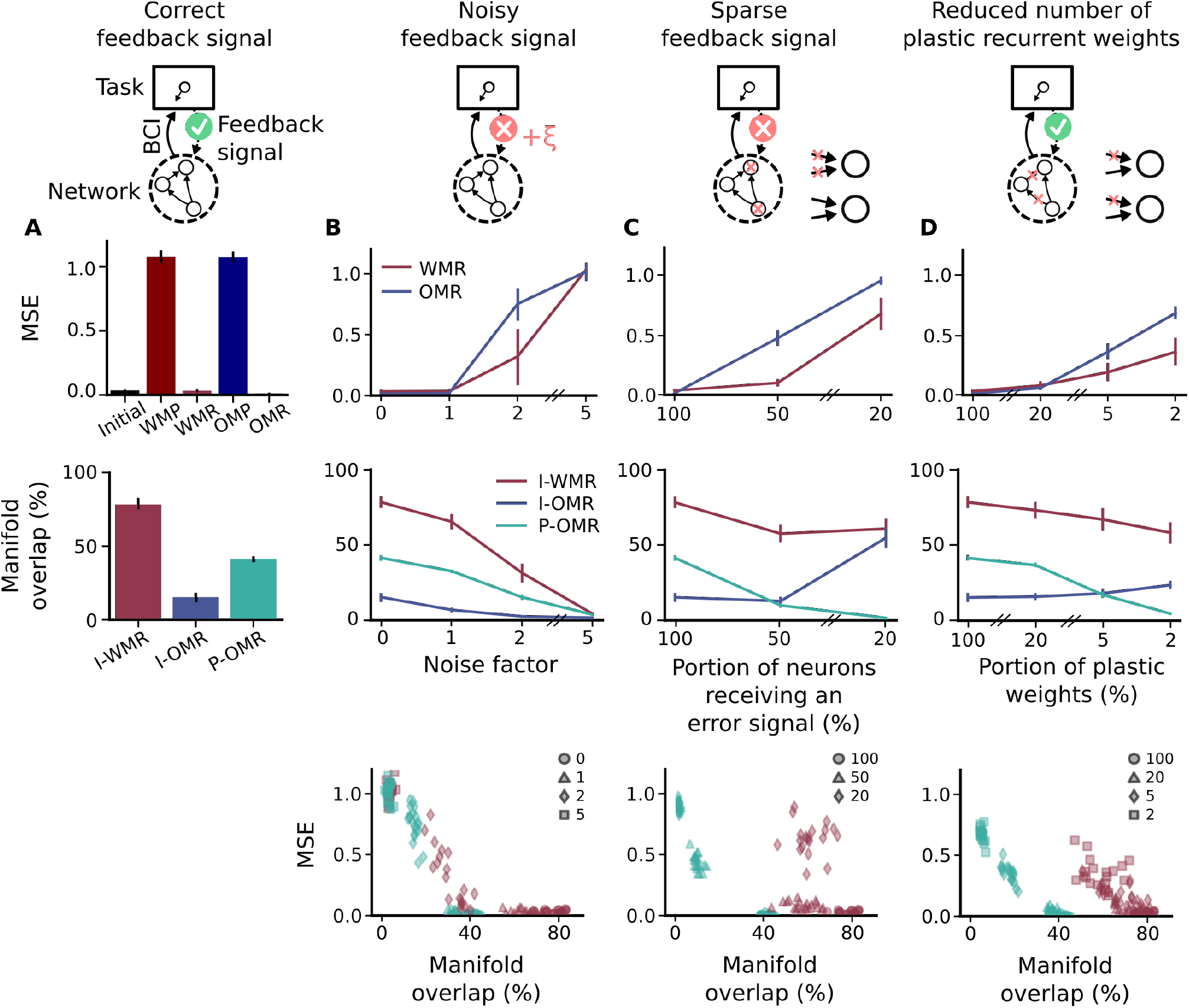
RNN with corrupted feedback signal or constraint on weight change can learn better within- than outside-manifold. (A) Relearning results for RNN with correct feedback signal (same as in Fig.2), bars indicate here and in the following standard deviation across networks. WMP and OMP show performance after within- or outside-manifold perturbations, whereas WMR and OMR show performance after retraining of the respective BCI perturbations (top). Manifold overlap is measured between initial manifold and manifold after within learning (I-WMR), initial manifold and manifold after outside learning (I-OMR) and manifold defined by BCI perturbation and manifold after outside learning (P-OMR) (center). (B) Relearning results for noisy feedback signal. The correct feedback transformation is distorted by adding independent noise to each entry of the matrix. The noise is drawn from a zero-mean Gaussian with standard deviation *σ* = *α* · *σ*_*original*_, where *α* is the noise factor and *σ*_*original*_ is the standard deviation of the correct feedback matrix. Top panel shows performance results, center panel shows manifold overlap and bottom panel shows the relation between both. (C) Relearning results for sparse feedback signal. In this scenario not all neurons in the recurrent network receive a feedback signal, which leads to a portion of recurrent weights remaining static during retraining. (D) Relearning results for sparse plastic connections. Similar to (C) a portion of recurrent weights remain static during retraining. But in contrast to (C), these static connections are not clustered to specific neurons. Instead, all neurons keep at least one plastic incoming connection.

Although the three different corrupted training scenarios show similar results with respect to retraining performance, the specific kind of disturbance could yet be highly different. Therefore, we analysed in more detail the network dynamics before and after retraining. We compared how well the internal manifold after retraining aligns with either the initial manifold in the case of within-learning, or with the dictated manifold coming form the BCI perturbation in the case of outside-learning. Interestingly, we found that, for noise in the feedback signal, the network dynamics are driven off the target manifold (Fig.3B center). This makes sense as the feedback signal is disturbed in random directions. During the retraining phase, the network then tries to align its internal manifold to these random directions, resulting in reduced overlap to the correct manifold dimensions. We found that task performance is high as long as manifold overlap is more than 40% (Fig.3B bottom), which is in line with what we found for outside manifold learning with the correct feedback signal (Fig.2D). In contrast, reducing the number of plastic connections has a different effect on recurrent restructuring. It primarily disturbs the manifold alignment for outside-manifold perturbations (Fig.3C & D center). This is due to the fact that the static connections in the network constrain the dynamics to stay within the original manifold. Therefore, for within-manifold learning the manifold overlap between before and after retraining is always relatively high (Fig.3C & D center). Nevertheless, the task performance is reduced in these cases as within-manifold perturbations can not correctly be counteracted (Fig.3C & D top). Furthermore, we found that not only the ratio of plastic connections affects retraining, but also the level of disturbance. As long as each neuron in the network receives a feedback signal, retraining performance is relatively high even if only 20% of the connections are plastic (Fig.3D top). In summary, although manifold overlap and task performance correlate in some regimes (e.g. Fig.3D bottom), we can find cases with high task performance and low manifold overlap (outside-manifold learning with correct feedback signal), and we can find cases with low task performance and high manifold overlap (within-manifold learning with sparse plastic connections), showing that one does not necessarily imply the other.

### Learning the feedback signal is possible for within- but not for outside-manifold perturbations

Although we have shown that within-manifold learning is more robust to corrupted feedback signals, the question remains of how the biological system could solve the error assignment and learn the feedback weights. A very naive hypothesis would be that it can infer the feed-back weights by simply knowing its own internal dynamics and regressing them against the observed cursor dynamics (Fig.4A). We tested this hypothesis and surprisingly found that this simple strategy gives good estimates of the feedback weights in the case of a within-manifold perturbation (Fig.4B), but fails completely in the case of an outside-manifold perturbation (Fig.4C). To quantify the quality of feedback learning, we calculated the correlation coefficient between the learned and the correct feedback weights. We found that the feedback learning performance is worse compared to inferring the feedback after the initial training phase for a within-manifold perturbation, but it is definitely better compared to an outside-manifold perturbation (Fig.4D). Furthermore, the estimate is almost as good after 6 trials as after 50 trials (Fig.4D). These results suggest that feedback learning is possible for within-manifold perturbations, at least from a computational point of view, and that observing only a few trials is in principle enough to obtain a good estimate. In contrast, the feedback weights can not be learned for an outside-manifold perturbation.

**Figure 4:**
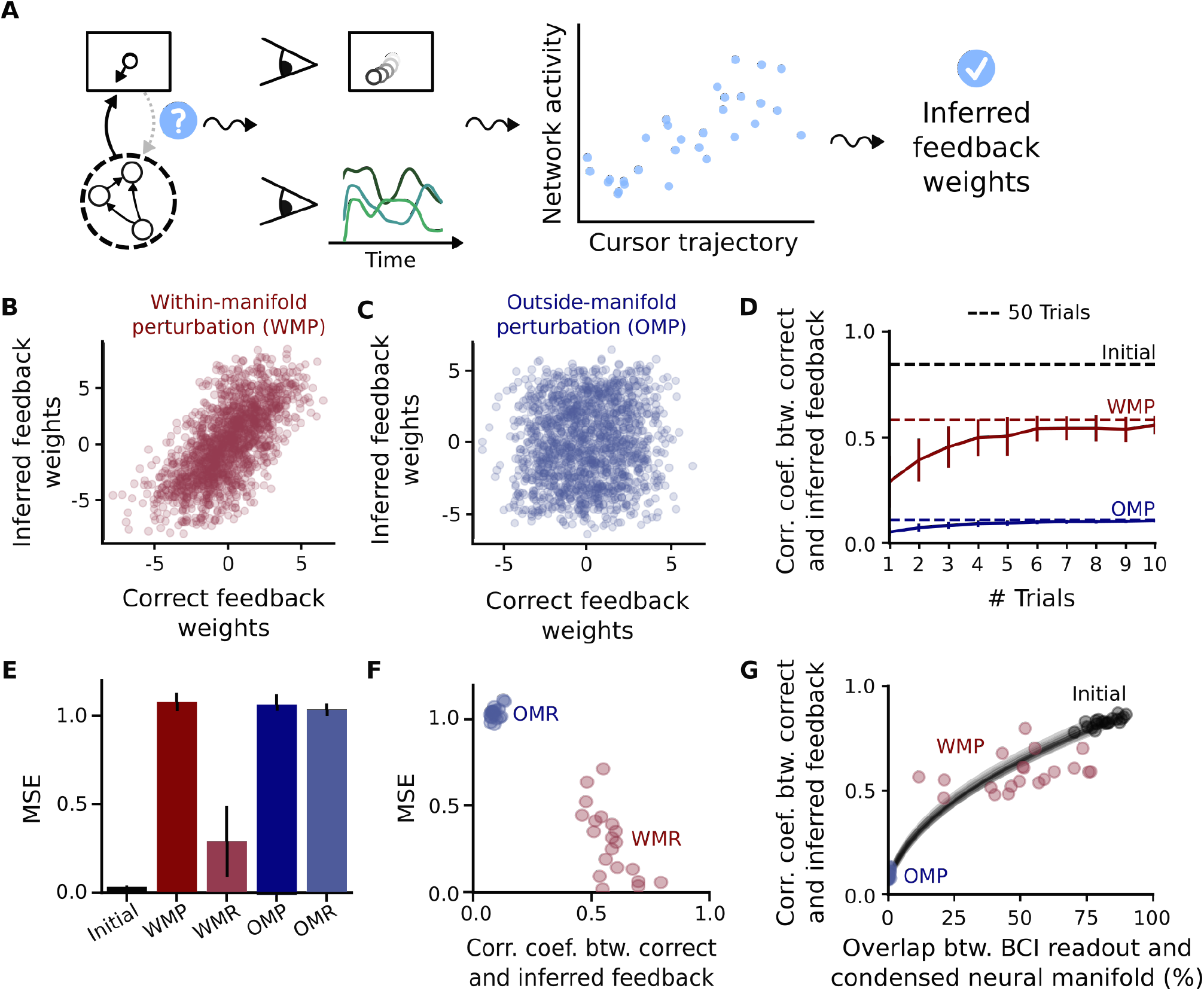
Learning the feedback signal is possible for within- but not for outside-manifold perturbations. (A) Regression is used to learn feedback weights. (B-C) Result of feedback learning for a within-manifold perturbation (WMP) (B) and an outside-manifold perturbation (OMP) (C). (D) Feedback learning results depend on the number of trials used for regression. Dashed lines show the feedback learning result for taking 50 trials. Compared is feedback learning after the initial training phase (where BCI readout and internal manifold completely overlap) (black), after within-manifold perturbation (red) and after outside-manifold perturbation (blue). (E) Recurrent relearning using the learned feedback weights. As before, task performance is quantified by calculating the mean squared error (MSE) between target cursor velocity and produced cursor velocity. Compared is task performance after within- (WMP) or outside-manifold (OMP) perturbation to performance after relearning with inferred feedback weights (WMR and OMR respectively) (F) Accuracy in the feedback learning, measured by the correlation coefficient between correct and inferred feedback weights, affects recurrent relearning performance, measured by mean squared error. (G) Alignment of internal manifold and readout determines feedback learning performance. To take into account variance in feedback learning for within-manifold perturbations, we calculate the manifold overlap only up to a network-specific dimension, not up to 10 dimensions as done in the rest of the paper. The specific dimension is defined as 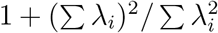 where λ_*i*_ are the Eigenvalues of the covariance matrix.

To get intuition for this result, we analysed a reduced model, where we synthesized data for any given Eigenvalue spectrum. For this reduced model, the output transformation only consists of the projection to the neural manifold, as this seemed to be the relevant part of the BCI mapping contrasting feedback learning for within- versus outside-manifold perturbations (Fig.S10). To learn feedback signals in this case, we regressed neural activity against 10-D neural mode activity (Fig.S11). By measuring feedback learning performance for different Eigenvalue spectra, we found that high dimensional dynamics would allow feedback learning even for outside-manifold projections (Fig.S12). To understand this, we revisited the definition of an outside-manifold perturbation. In the case of an outside-manifold perturbation, shuffling creates a new set of ‘wrong’ Eigenvectors, where each vector can be described as a superposition of true Eigenvectors of the system. For low-dimensional dynamics, there are only a few highly prominent modes, which consequently also dominate the produced mode dynamics obtained with the ‘wrong’ Eigenvectors, irrespective of the specific weight in the superposition. Our regression algorithm, used to infer feedback weights, is biased towards these highly dominant modes, which leads in the end to incorrectly inferred feedback weights. In fact, the inferred feedback weights for an outside-manifold perturbation in our main simulation setup always pointed in directions which are within the original manifold. This seemingly counter-intuitive result might be explained by the aforementioned reasoning.

In a next step, we wanted to test whether the learned feedback weights are good enough to correctly drive recurrent relearning. For this, we estimated the feedback weights offline and then used them during the recurrent relearning period. As expected, we found that outside-manifold learning will be completely impaired if we use the learned feedback weights (Fig.4E blue). In contrast, task performance increased during recurrent relearning for within-manifold perturbations (Fig.4E red). Interestingly, we found that there is a relatively large amount of variation in relearning performance if we compare different network initializations. However, this large amount of variation is only observed for the task performance after within-manifold relearning. We hypothesized that the accuracy of the learned feedback weights is responsible for the successful retraining of recurrent weights during the adaptation phase. This is indeed what we found when we plotted the retraining performance against the feedback learning accuracy (Fig.4F). In summary, these results suggest that the accuracy of the feedback learning determines whether or not the BCI perturbation can be counteracted by recurrent restructuring. Together with the fact that it is not possible to infer feedback weights for outside-manifold perturbations, this could explain why outside-manifold learning was not observed experimentally [Sadtler et al., 2014].

Although we have shown that the accuracy of the feedback signal is responsible for either success or failure of recurrent retraining, it is still unclear why feedback learning works better for some BCI perturbations than others, and especially why it works for within-manifold perturbations, but completely fails for outside-manifold perturbations. As the main difference between within- and outside-manifold perturbations is the alignment of the readout to the internal manifold, we hypothesized that this alignment controls whether feedback learning is successful or not. To test this, we carried out simulations where we interpolated between the initial BCI readout and the one used for an outside-manifold perturbation. This allowed us to sweep the whole spectrum of alignments, as the initial BCI readout is completely aligned to the internal manifold of the network, whereas the readout for outside-manifold perturbations is completely orthogonal to it. We then inferred the feedback weights for all of these different readouts and quantified the result by calculating the correlation coefficient between the learned and the correct feedback weights. As hypothesized, we found that there is a specific monotonic relationship between feedback learning performance and alignment of BCI readout and internal manifold (Fig.4G). This also explains why we observed a large variation for within-manifold relearning performance. In our setup, we fix the number of manifold dimensions used in the BCI readout, but for each individual network the internal manifold dimension can deviate from this. This means that within-manifold perturbations can be in fact slightly outside-manifold for individual networks, so that the BCI readout in these cases is not hundred percent aligned to the true internal manifold. In summary, by looking at the underlying requisites enabling successful feedback learning, we could understand why it is impossible to infer correct feedback weights in the case of an outside-manifold perturbation and why we would expect variations in within-manifold learning in the case that the readout dimension does not exactly match the internal manifold dimension.

### Feedback signal for outside-manifold perturbation can potentially be learned with incremental strategy

Besides the original experimental study showing that monkeys can not adapt to outside-manifold perturbations on the timescale of a single experiment [Sadtler et al., 2014], there is new evidence showing that outside-manifold learning is indeed possible if the training lasts for several days and follows a specific incremental protocol [Oby et al., 2019]. We have shown so far that outside-manifold learning should not be possible, as the correct feedback weights needed for successful retraining can not be learned. We reasoned that to nevertheless be able to learn outside of the original manifold, one would have to insert intermediate perturbation steps, where for each perturbation step there is a certain amount of overlap between the current internal manifold and the BCI readout. This would assure that the system has the chance to learn constructive feedback weights in each step, which could then lead to successful recurrent restructuring. With this strategy, one could potentially build up the learning by shifting the internal manifold step-by-step closer to the one defined by the outside-manifold perturbation. To test this idea we implemented the incremental strategy by inserting three intermediate perturbations which interpolate between the original BCI readout and the readout for an outside-manifold perturbation (Fig.5A).

**Figure 5:**
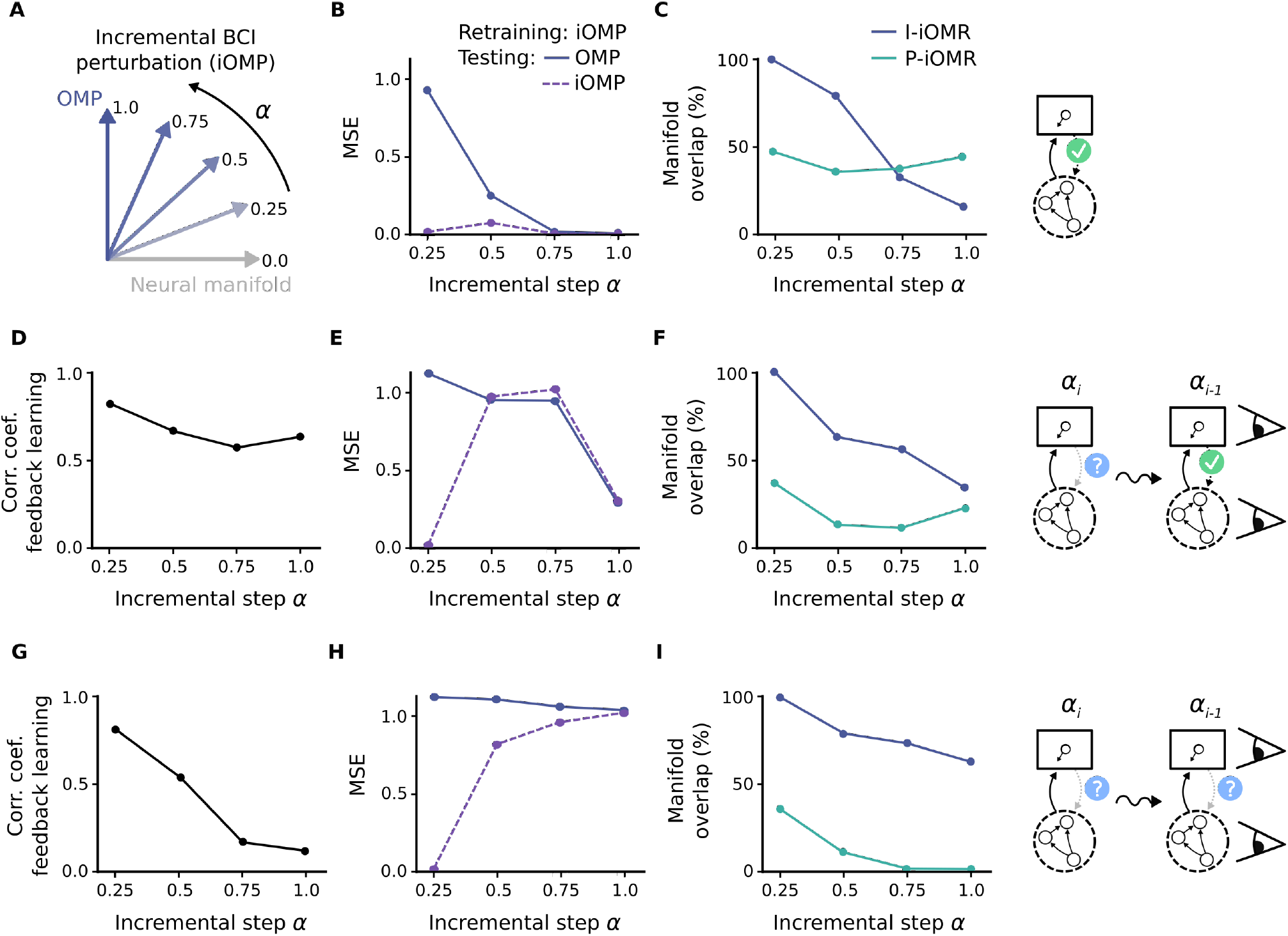
Feedback signal for outside-manifold perturbation can potentially be learned with incremental strategy. (A) Incremental outside-manifold perturbation, where with each incremental step *α* the BCI readout is less aligned with the original neural manifold. (B-C) Retraining results for incremental perturbation steps using correct feedback signal. (B) Task performance after retraining, measured as mean squared error (MSE), achieved with current incremental BCI readout (iOMP), which is the readout used during retraining, as well as achieved under the full outside-manifold perturbation (OMP). (C) Manifold overlap between initial manifold and manifold after relearning (blue line), as well as between target manifold, defined by the current BCI readout, and manifold after relearning (green line). (D-F) Retraining results using incremental strategy and assuming, that each consecutive incremental learning step starts from a correctly trained network. (D) Feedback learning performance, quantified by calculating the correlation coefficient between correct and inferred feedback weights, for each incremental learning step. (E) Task performance after recurrent retraining. (F) Manifold overlap for each incremental step. (G-I) Retraining results using full incremental strategy. Here, each consecutive learning step starts from the network state obtained in the previous step. Feedback learning performance (G), task performance after recurrent retraining (H) and manifold overlap (I).

To test whether the inclusion of the extra BCI perturbations is enough to enable feedback learning, we firstly performed a simplified simulation, where the feedback learning relies on a correctly trained previous network state (Fig.5D-F). To obtain the correctly trained network state, we ran a separate retraining simulation with an ideal-observer feedback signal for every intermediate perturbation step. As expected, each of the incremental BCI perturbations could correctly be counteracted by recurrent retraining with an ideal-observer feedback signal (Fig.5B). By looking again at the manifold overlap obtained during retraining, we found that each retraining simulation achieved between 40% - 50%, in line with our previous results (Fig.5C). To test feedback learning, we next tried to learn the correct feedback weights for each perturbation step by observing the correctly trained network from the preceding perturbation step. We found that the overlap between the network state of one perturbation step and the BCI readout of the next step is high enough to allow successful feedback learning (Fig.5D). To finally test whether the incremental learning strategy would work in this simplified setup, we used the learned feedback weights to retrain the recurrent network. Interestingly, we found that retraining drastically improves by using an incremental training strategy (Fig.5E). However, the manifold overlap achieved during recurrent retraining is lower compared to retraining with an ideal-observer feedback signal (Fig.5E). This makes sense as the learned feedback signal does not completely point in the new desired directions, as it would do for the correct feedback signal. Instead, it also has components pointing in the dimensions of the manifold orientation before retraining. With this simplified scenario we could show that incremental learning can potentially be used to counteract an outside-manifold perturbation, assuming that the new incremental perturbation always starts from a well-trained network.

To finally see whether our network can be trained incrementally, without relying on correctly trained intermediate steps, we implemented a similar simulation as before, but now the feedback learning uses the preceding network states which also have been trained with learned feedback, which is not necessarily the correct feedback we have used before (Fig.5G-I). We found that for this scenario the accuracy of the feedback learning drops for each intermediate perturbation step (Fig.5G). As expected, this deficiency led to poor performance in the recurrent retraining (Fig.5H) and the incremental strategy is therefore not helping in this case. To understand why incremental learning fails here, we compared what happens to the overlap between internal manifold and BCI readout with each consecutive step. In the simplified simulations before (Fig.5B-F), we found that the internal manifold does not align as well with the current readout if we use learned feedback weights instead of the correct ones (compare Fig.5C and Fig.5F). In the following incremental step, this reduced overlap leads to a larger misalignment between current network state and BCI readout, which in turn leads to a decrease in feedback learning accuracy. The problem is therefore that the network can not catch up with the BCI perturbation and falls behind more and more for every incremental learning step. In summary, we demonstrated that a progressive schedule could in principle lead to outside manifold learning, but we also identified limitations associated with this schedule.

## Discussion

To investigate which factors influence learning within and outside of the neural manifold, we used a recurrent neural network trained with a machine learning algorithm. Although this method of modelling neural dynamics lacks biological details, using RNNs has been surprisingly useful to understand neural phenomena [Sussillo et al., 2015, Barak, 2017, Mastrogiuseppe and Ostojic, 2018, Michaels et al., 2019, Masse et al., 2019]. Using this approach, we could identify feedback learning as a potential bottleneck differentiating between learning within versus outside the original manifold. As feedback learning might not be the only relevant factor, we also tested the impact of further biological constraints. We found that neither separation of excitatory and inhibitory neural population (Fig.S5), nor local learning rule (Fig.S6) [Murray, 2019, Bellec et al., 2019], qualitatively change the fact that within- and outside-manifold perturbations can equally well be learned, given an ideal-observer feedback signal.

In principle, there are two alternative scenarios which would lead to altered network dynamics in motor cortex. Assuming motor cortex is highly controlled by other brain regions, such as for example the cerebellum and pre-motor cortex, the first scenario would be to alter the control signals coming from these regions, and to therefore trigger different activity patterns [Menendez and Latham, 2019]. Nevertheless, how the inputs are learned remains unclear. The second scenario is rewiring in the local circuit. In this study we focused on the latter and tested to what extent it can explain experimental observations [Sadtler et al., 2014, Oby et al., 2019]. We found that reorganization of local connectivity does not necessarily lead to a change in neural manifold. This shows that globally coordinated changes in neural activity can not only be realized by adaptation of input drive, but also via local rewiring driven by a learned feedback signal. It remains to be investigated to what extent one can contrast both frameworks, given experimental data, as biology potentially uses multiple mechanisms in parallel.

A recent computational study has shown that weight changes necessary for within-manifold learning are much smaller compared to outside-manifold learning and proposed this finding as a potential reason for the experimental observation in Sadtler et al. [Wärnberg and Kumar, 2019]. In contrast, we could not observe a difference in weight change during retraining (Fig.2C). However, we did observe that, if we constrain the number of plastic weights, within-manifold learning is more successful compared to outside-manifold learning (Fig.3C-D). One fundamental difference compared to Wärnberg and Kumar is that, in our study, the weight dynamics are not constrained to one dimension. Instead, by calculating the effective rank of the weight change during relearning we found that weight dynamics explore various dimensions (see Fig.S4). Having a more flexible training algorithm, which is potentially able to explore all weight configurations, could explain why we found that small weight adaptations are sufficient for successful within- as well as outside-manifold learning. Which one of these two scenarios is more plausible in a biological context is unclear. Thus, any conclusions drawn critically depend on further investigation of biological learning dynamics.

As having a feedback signal is a general concept of motor control [Kawato, 1999, Wolpert and Ghahramani, 2000, Todorov and Jordan, 2002], it would be interesting to study whether one could also infer correct feedback signals in an upstream region exclusively for the case of a within-manifold perturbation. We hypothesize that, as long as the, potentially non-linear, output transformation via motor cortex is aligned with the internal manifold of the upstream region, feedback learning should be viable. Thus, our study gives a new perspective on the general problem of credit assignment, proposing that correct feedback learning critically depends on the alignment between output transformation and internal manifold in a specific neural pathway.

Our results can also be related to a recent finding, reporting that a variety of different movements are controlled by neural activity lying in the same subspace across movements [Gallego et al., 2018]. As our results suggest that the readout - used to produce a movement - indirectly defines the accessible neural activity space for motor learning (Fig.4F & G), we would expect that a stable readout implies a stable manifold across different movements. One could test this hypothesis by estimating the muscle output transformation for each specific movement. We would predict that less overlap in the output transformation implies less overlap in neural manifolds, measured during the two different movements.

Finally, our work might be relevant for future BCI designs. Our results support the idea that a subject can potentially learn to interact with any kind of static BCI mapping, as long as its dimensions are aligned to the internal dynamics of the local circuit it is connected to. It remains to be experimentally tested to what extent this holds true, but if so, it could impact development of brain-computer interfaces for real-world applications.

In summary, our model makes the following predictions, 1) for within-manifold perturbations, it is possible to infer useful feedback weights by regressing neural activity against cursor dynamics, 2) learning performance during a BCI task can be predicted by the alignment between BCI readout and internal manifold of the connected brain region, 3) silencing local network rewiring would impair learning of within- as well as outside-manifold perturbations.

In conclusion, we showed that local network rewiring can account for the observed behavioural differences between within- and outside-manifold learning, assuming that the feedback signal driving local reorganization has to be learned.

## Methods

**Table 1:**
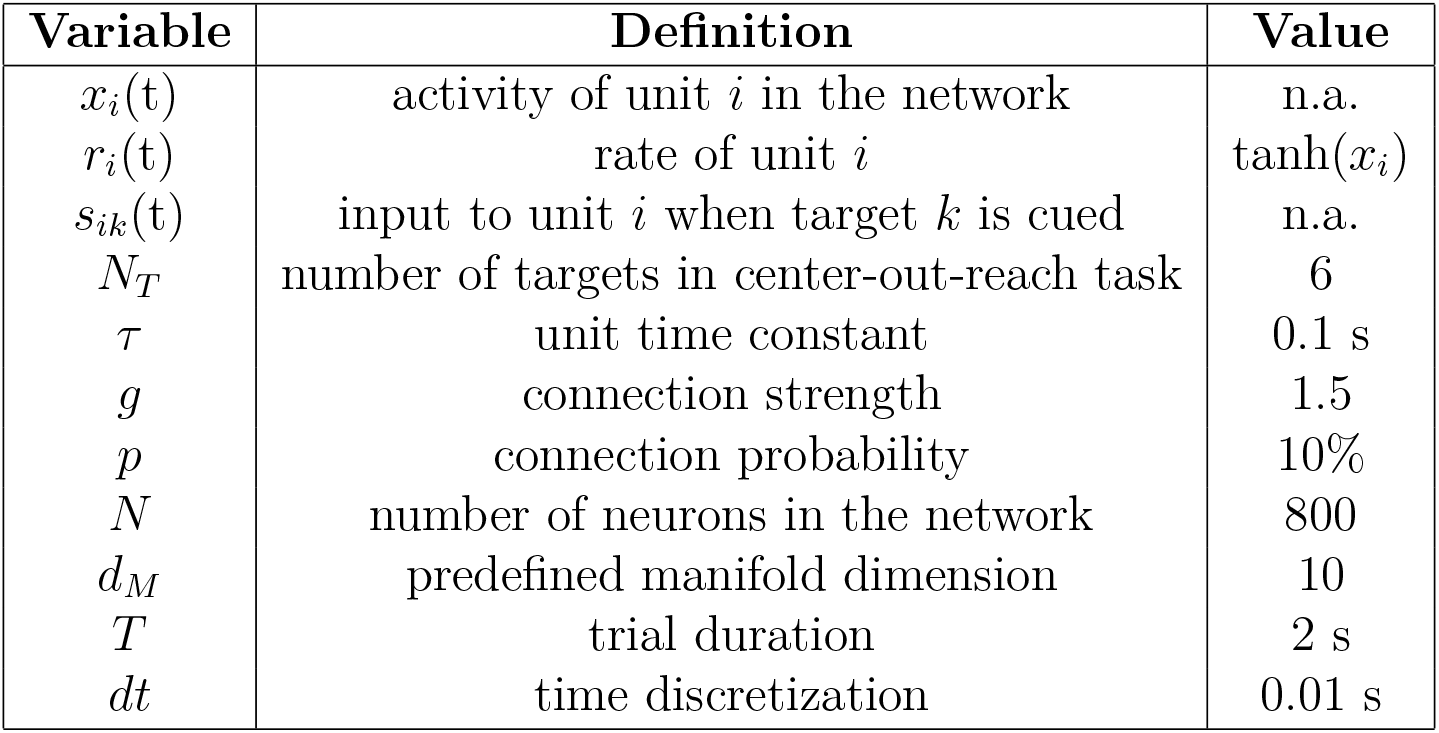
List of parameters used in the model.

### Task

In order to compare our learning results to experimental studies, we implemented a standard center-out-reach task, which is regularly used to experimentally probe motor learning. In our case, the task consists of six targets, evenly distributed on a circle. In an experiment, the subject would get a visual cue for example at the beginning of each trial to indicate which of the six targets is the target of the current trial. The subject should then reach to this target in a given amount of time. We simulated the target cue by having an input pulse at the beginning of each trial, which is distinct for each of the six different targets. To simulate activity in motor cortex we use a recurrent neural network (RNN). The output is produced by a mapping, which we called brain-computer interface (BCI) for didactic reasons. It transforms the rates measured in the RNN to *x* and *y* velocity of a cursor, where the cursor simulates the center-out-reach movement. At the beginning of each trial, the cursor is set to the center of the workspace, and after the target cue ends, its movement is controlled by the RNN dynamics. The target speed for the cursor is 0.2 and the target directions are evenly distributed between 0 and 2*π*. To measure performance, as well as for training, we calculated the mean squared error between the target cursor velocity and the produced cursor velocity. As the target cursor velocity is constant and does not change depending on where the cursor is at a given moment, we use an open loop setup. The target for each specific trial is randomly chosen among the six.

### Recurrent neural network (RNN)

The recurrent neural network is simulated by the following dynamical equation:

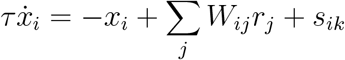

The recurrent connections *W*_*ij*_ are sparse, with connection probability *p* = 10%, and drawn from a Gaussian 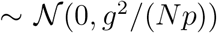. To simulate the target cue our network receives a pulsed input *s*_*ik*_ at the beginning of each trial. The input is given by

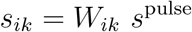

where *s*^pulse^ is a 0.2*s* long pulse with amplitude 1 and the input weights *W*_*ik*_ are drawn randomly from a uniform distribution 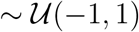.

To implement learning in the network we use an adapted form of the recursive-least-squares algorithm, which is inspired from [Laje and Buonomano, 2013]. If learning is on, it starts after the target cue period. Then, at every second time point (which corresponds to every 20*ms*) an update step is made. Firstly, the performance error *e*^*P*^, given by the mean squared error between the target cursor velocity and the produced cursor velocity, is calculated. Via a linear transformation this performance error is then assigned to errors *e* on the activity of single units in the network.

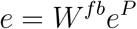

This transformation *W*^*fb*^ is what we define as feedback weights. In the ideal-observer case, this feedback matrix is given by the pseudo-inverse of the BCI transformation. Note that we calculate the error for each neuron through these feedback weights, which is used to adapt recurrent weights in the network, but there is no additional input for each neuron due to this error signal. The recurrent weight update is given by

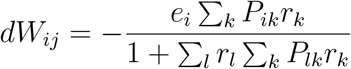

where *e*_*i*_ is the error for unit *i* and *P* is a matrix which estimates the inverse of the correlation matrix. *P* is also updated in every update step and the update rule is, in matrix form, given by

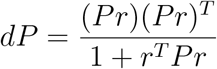

At the start of learning, the matrix *P* is initialized as diagonal matrix with 0.05 on the diagonal. For each training period the network is trained for 80 trials. Note that the training algorithm does not change the structural connectivity, so only connections which are non-zero at the beginning are updated.

### Brain-computer interface (BCI)

The brain-computer interface consists of two linear transformations which sequentially transform network rates to cursor velocities. The first transformation is given by a projection to principle components of the network dynamics. To calculate this projection, we measured the dynamics over 50 trials during which the network is performing the task. Here, and at every other point, we took network dynamics into account only from the end of the cue phase. Then, we used principle component analysis to calculate the projection matrix *C*. The second transformation is given by a decoder matrix *D* which is optimized offline by doing regression of target cursor velocities against the first 10 principle components, taking the same 50 trials as before. The full BCI transformation *T* is then given by

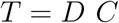

For the initial training period, *T* is set up randomly. Each entry of the matrix is drawn from a standard normal distribution, normalized to have matrix norm 1 and multiplied with a factor 0.04. This setup showed to drive initial learning sufficiently well, and assured good task performance after the initial network training period, followed by the BCI tuning described above (Fig.S2). Note that the initial random mapping has high impact on the produced neural manifold and therefore also on the optimized BCI mapping (Fig.S9).

For a within-manifold perturbation, we insert a permutation matrix *η*_*WM*_ in between the first and the second linear transformation, which shuffles the decoder weights related to the principle components.

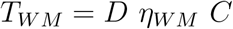

In contrast, for an outside-manifold perturbation, we insert a permutation matrix *η*_*OM*_ before the first linear transformation, which shuffles the projection matrix and therefore leads to not extracting the correct principle components of the network dynamics.

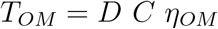

In order to assure that both types of perturbations cause approximately the same performance error we calculate the performance for 200 random perturbations of each type and then pick the ones which closely match in the performance error they cause.

### Analysis

To quantify manifold overlap we adapted the measurement used in [Golub et al., 2018]. Firstly, we calculated the covariance matrix of the network dynamics *S*, given 50 trials of performing the task. This was done for the initial network (subscript _1_) and for the network after retraining (subscript _2_). Then, both of these matrices were projected onto the neural manifold defined by the initial network. This projection is given by the same projection matrix *C* as used in the BCI transformation. To quantify the amount of explained variance in the first 10 dimensions, defined as *β*_1_ for the initial network and *β*_2_ for the retrained network, we calculated the trace of the projected matrices (taking into account only the first 10 dimensions) and divided it by the trace of the original matrices (full-dimensional).

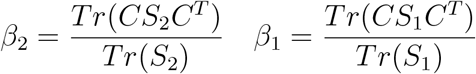

The manifold overlap is then defined as the ratio *β*_2_/*β*_1_. For calculating the overlap between the perturbed BCI readout and the network manifold after retraining we replaced the projection into the Eigenbasis of the initial covariance matrix, given by *C*, with a projection given by the new readout transformation, which is *C η*_*OM*_.

To calculate the overlap between BCI readout and condensed network manifold (Fig.4G), we summed up the normalized projections of the readout *T* into the first *k* dimensions of the Eigenspace of the network dynamics, given by *C*,

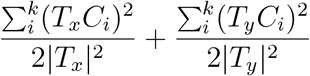

where *T*_*x*_ is the first row of the BCI transformation, decoding the cursor velocity in x-direction, and *T*_*y*_ is the second row of the BCI transformation, decoding the cursor velocity in y-direction. The number of dimensions *k* is defined by

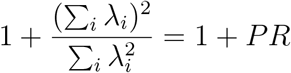

where λ_*i*_ are the Eigenvalues of *S*_1_ and *PR* is participation ratio, a commonly used measure of dimensionality.

### Feedback learning (Fig.4)

To learn the feedback weights for a given BCI perturbation, we created a dataset of 50 trials where the network is performing the task with the perturbed BCI readout. During this, there are no plastic changes yet in the recurrent connections. To infer feedback weights we then regressed the network rates, measured during the 50 trials, against the x and y cursor velocities. The accuracy of the regression is about the same for within- and outside-manifold perturbations, whereas the match between inferred and true feedback weights strongly differs. To obtain Fig.4D, we only used a randomly selected subset of the 50 trials. To obtain Fig.4G, we analysed not only the given BCI perturbations, but added intermediate perturbations, which interpolate between the original BCI transformations and the outside manifold perturbations. The analysis of the interpolated perturbations was shown in Fig.4G (black lines).

### Incremental strategy (Fig.5)

In each incremental step, the learning starts from the weight configuration after initial training. We did this to insure that the network stays in a physiological state, as the recursive-least-squares algorithm tends to increase the weights during training. Without this reset, the weight distribution would grow with every incremental training and could potentially lead to a network state where most of the neurons are saturated.

Each incremental BCI perturbation *T*_*α*_ is defined by interpolating between the original readout *T* and the full outside-manifold perturbation *T*_*outside*_, where the incremental factor *α* defines the ratio between both.

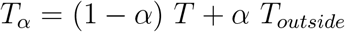

## Acknowledgements

We thank A. Kumar, J.A. Hennig, S.M. Chase, B.M. Yu, A.P. Batista for discussions and J.A. Gallego and E.R. Oby for discussions and comments on the manuscript. This work was supported by BBSRC BB/N013956/1, BB/N019008/1, Wellcome Trust 200790/Z/16/Z, Simons Foundation 564408 and EPSRC EP/R035806/1.

