## Supplemental Figures for "Neural manifold under plasticity in a goal driven learning behaviour"

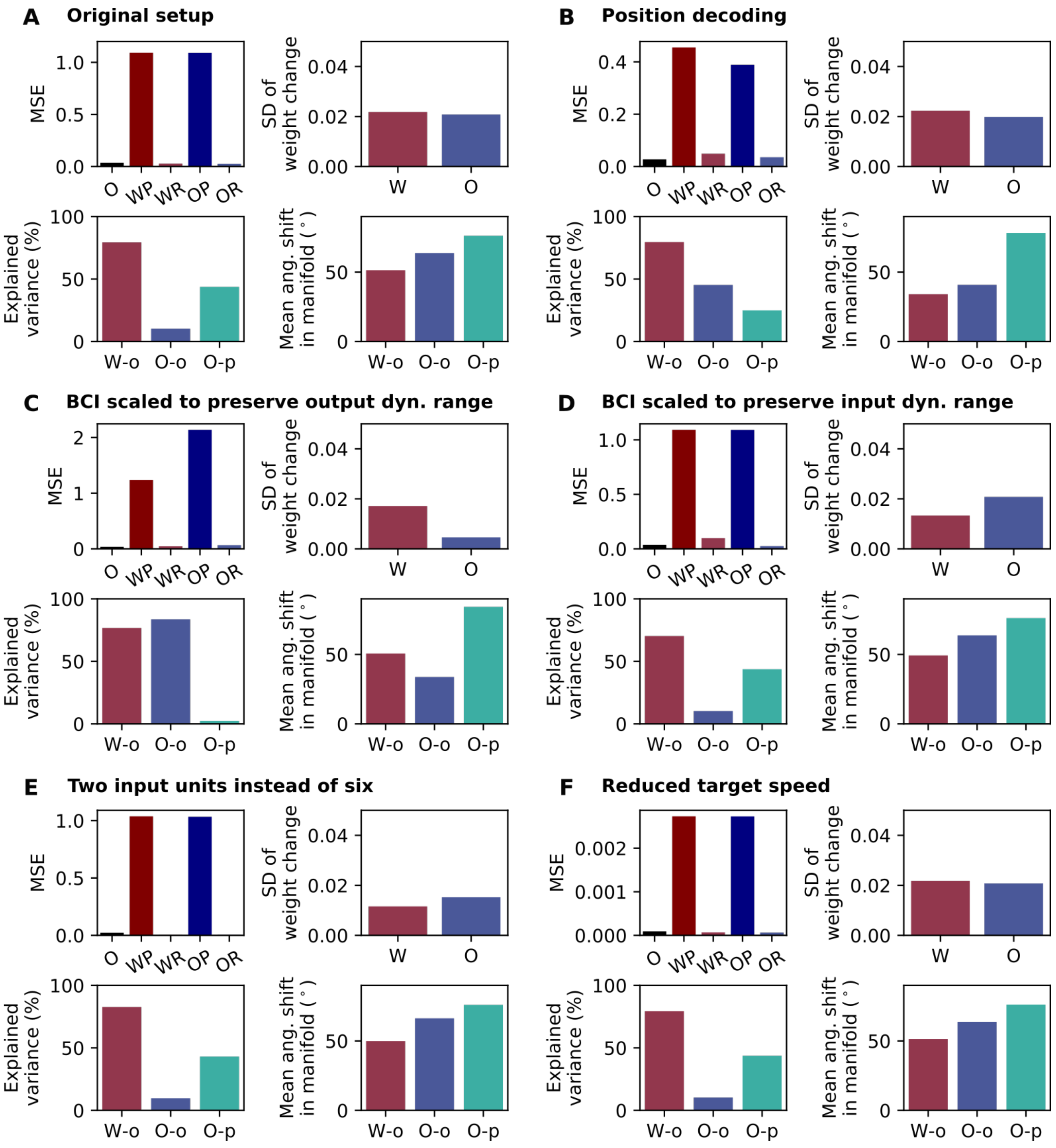


**Figure S1: Control simulations.** (A) Original setup as in Fig.2. Upper left panel shows task performance, measured as mean squared error (MSE), after initial training (O), after within-manifold perturbation (WP), after within-manifold retraining (WR), after outside-manifold perturbation (OP) and after outside-manifold retraining (OR). Upper right panel shows the standard deviation of the weight change distribution between before and after retraining for within- (W) and outside-manifold (O) retraining. Lower left panel shows the manifold overlap between initial and retrained manifold for within- (W-o) and outside-manifold (O-o) perturbation, as well as the overlap between retrained and target manifold for outside-manifold perturbations (O-p). Lower right panel shows the mean principle angle between the same manifold as in lower left panel. (B) Position decoding instead of velocity decoding. (C) BCI perturbations are scaled in order to preserve the range of velocities after perturbation. This especially affects outside-manifold perturbations as in this case the velocity values are normally only 10% of the original ones, due to a signal loss caused by the outside-manifold projection. (D) BCI perturbations are scaled to assure that the theoretical approximation for the neural activity after retraining, given the initial training state, does not exceed the dynamic range of the neurons. (E) Instead of having one input unit for each target, here, there are only two input units. They signal x and y position of the targets. (F) Reduced target speed of 0.01, instead of 0.2.

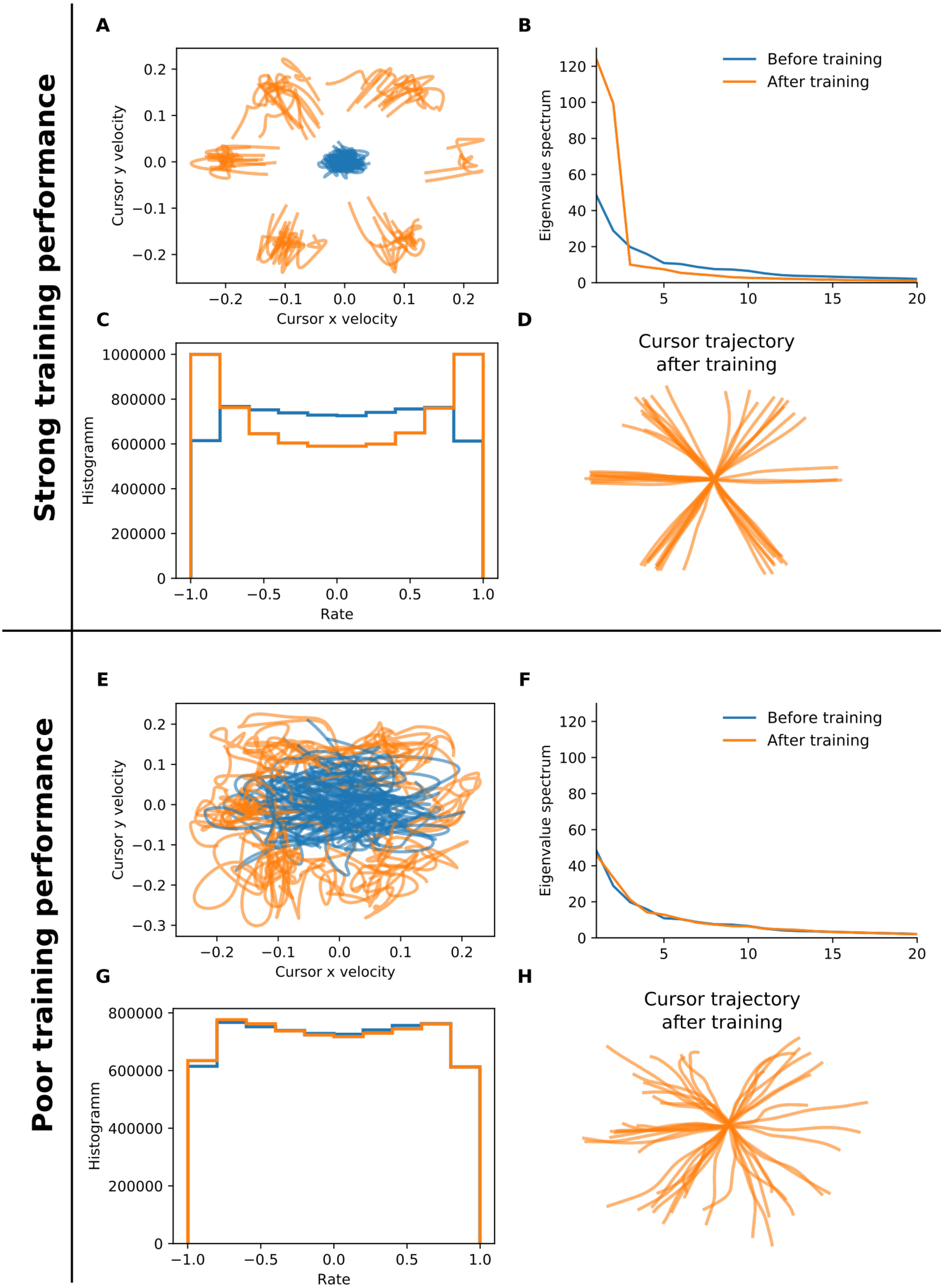


**Figure S2: Details of initial network training.** (A-D) Initial training with random decoder, normalized and scaled with factor 0.04 (the default version used in the paper). (E-H) Initial training with random decoder, normalized and scaled by a factor 0.2. (A)&(E) Cursor velocities before and after training. (B)&(F) Eigenvalue spectrum of network dynamics before and after training. (C)&(G) Rate distribution before and after training. (D)&(H) Reconstructed cursor trajectory after training.

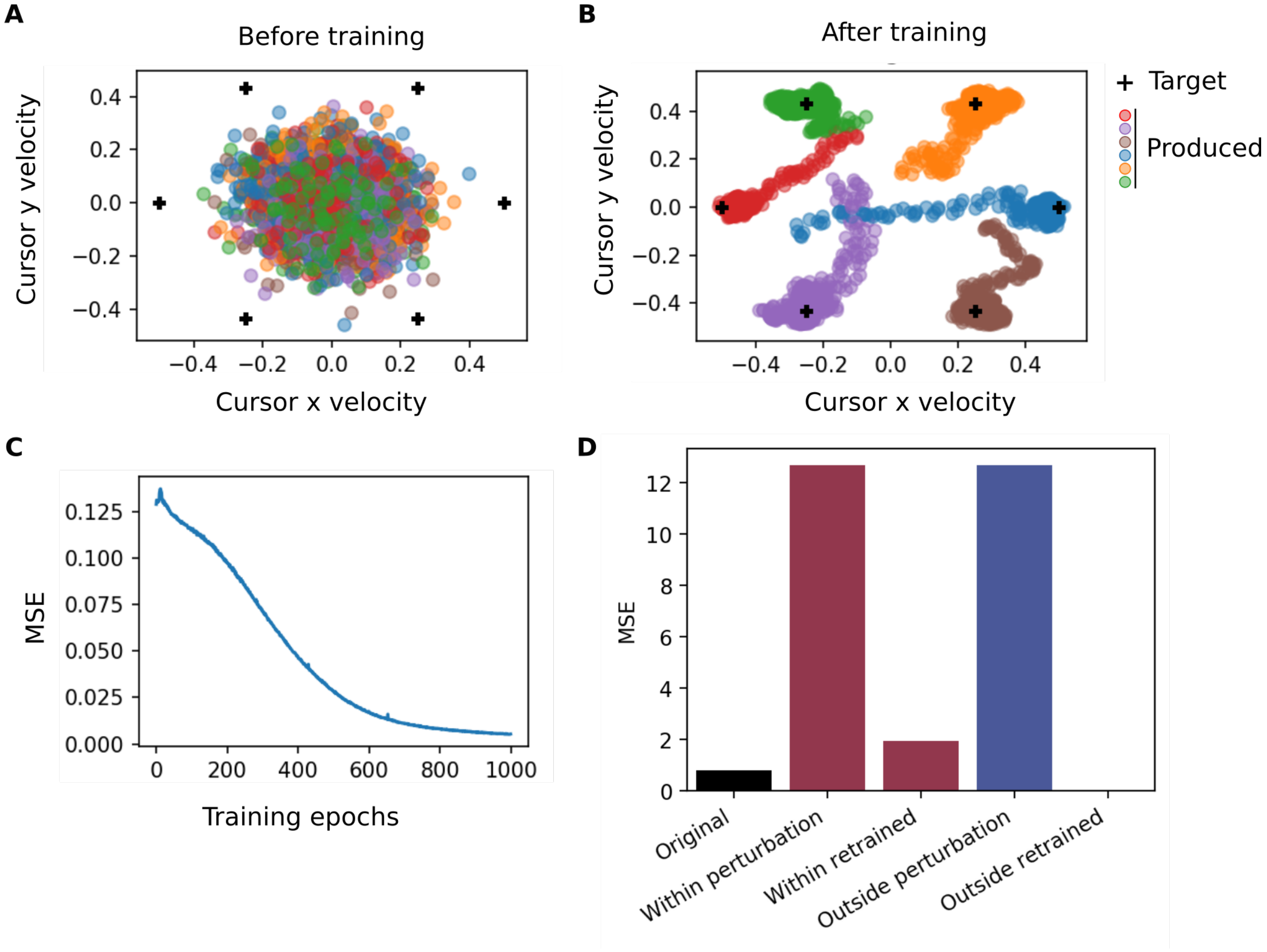


**Figure S3: Simulation results for recurrent neural network trained with backpropagation-through-time using Pytorch.** Here, the network dynamics are given by $h_t = \tanh(W_{ih}x_t + W_{hh}h_{(h-1)})$. Each trial has 100 time steps and the target cue is given during the first 5 steps. The input $x_t$ is modelled similar to our main setup. The stimulus amplitude is 50. The incoming weights $W_{ih}$ are fixed and randomly drawn from a uniform distribution between $-\sqrt{1/N}$ and $\sqrt{1/N}$, where $N$ is the number of neurons in the network. The recurrent weight matrix is initialized in a similar fashion as in our main setup, except that here, we use a fully connected network. For gradient descent we use Adam optimizer [Kingma and Ba, 2014] with learning rate 0.001. The loss and the fixed output decoder is the same as in our main setup. The training uses an ideal-observer feedback signal to propagate the error in cursor velocities to errors on single neurons in the recurrent network. For training, we use a batch size of 30. (A) Cursor velocities before initial training. (B) Cursor velocities after initial training. (C) Learning curve for initial training. (D) Performance results for within- and outside-manifold retraining. Number of training epochs for initial training, as well as retraining, is 1000.

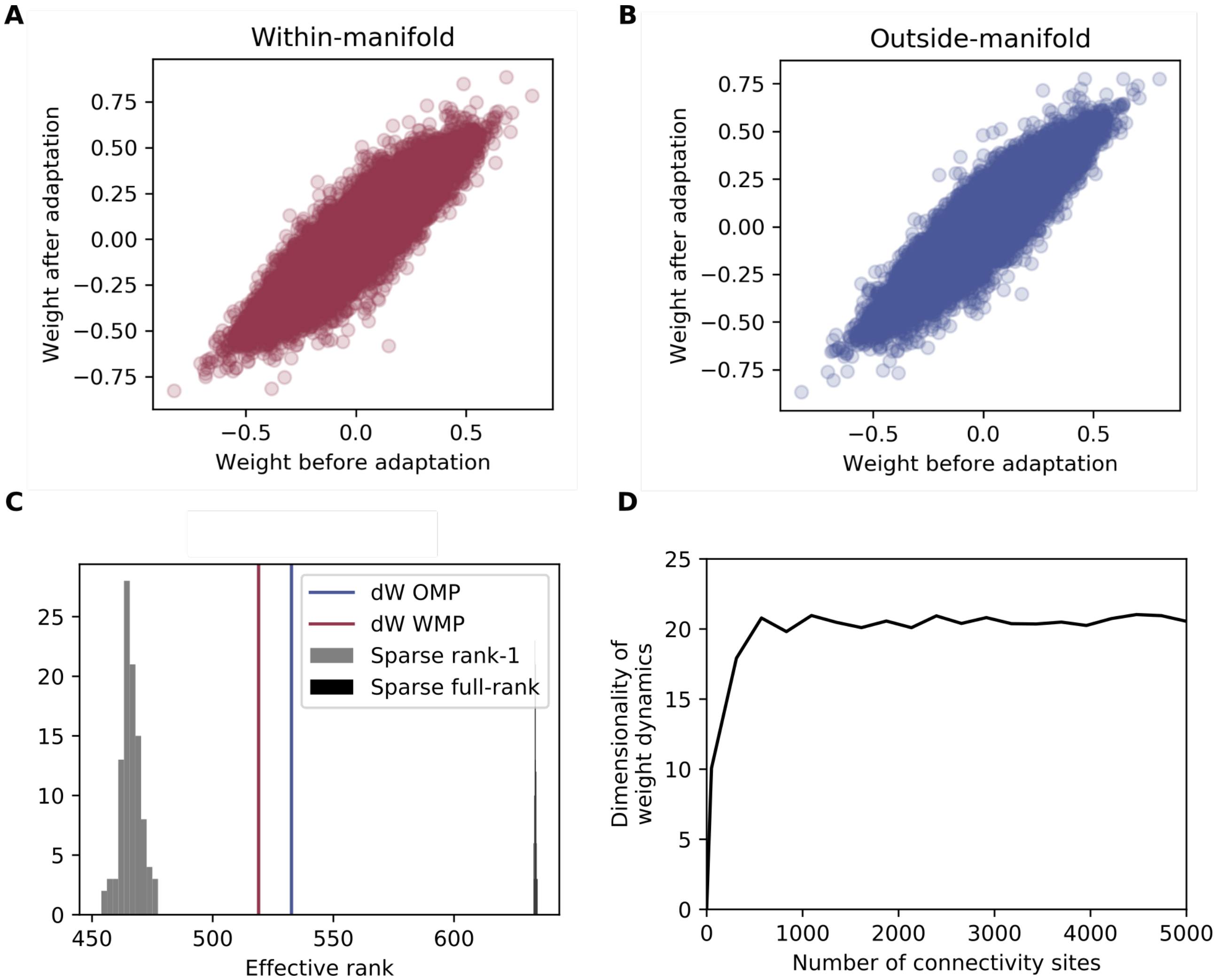


**Figure S4: Weight change during retraining.** (A) Relation between weight before retraining and after retraining for within-manifold perturbation. (B) Relation between weight before retraining and after retraining for outside-manifold perturbation. (C) Measurement of the effect rank [Roy and Vetterli, 2007] of the weight change matrix for within-manifold and outside-manifold retraining. Both values are compared to values obtained for random rank-1 and full-rank matrices, having the same sparsity as the network weight matrix. (D) Dimensionality of weight change dynamics during retraining. As it is not feasible to calculate the full covariance matrix for all plastic connections, as the number is too high ($\sim 64000$), we calculated the dimensionality from randomly chosen subsets of connections.

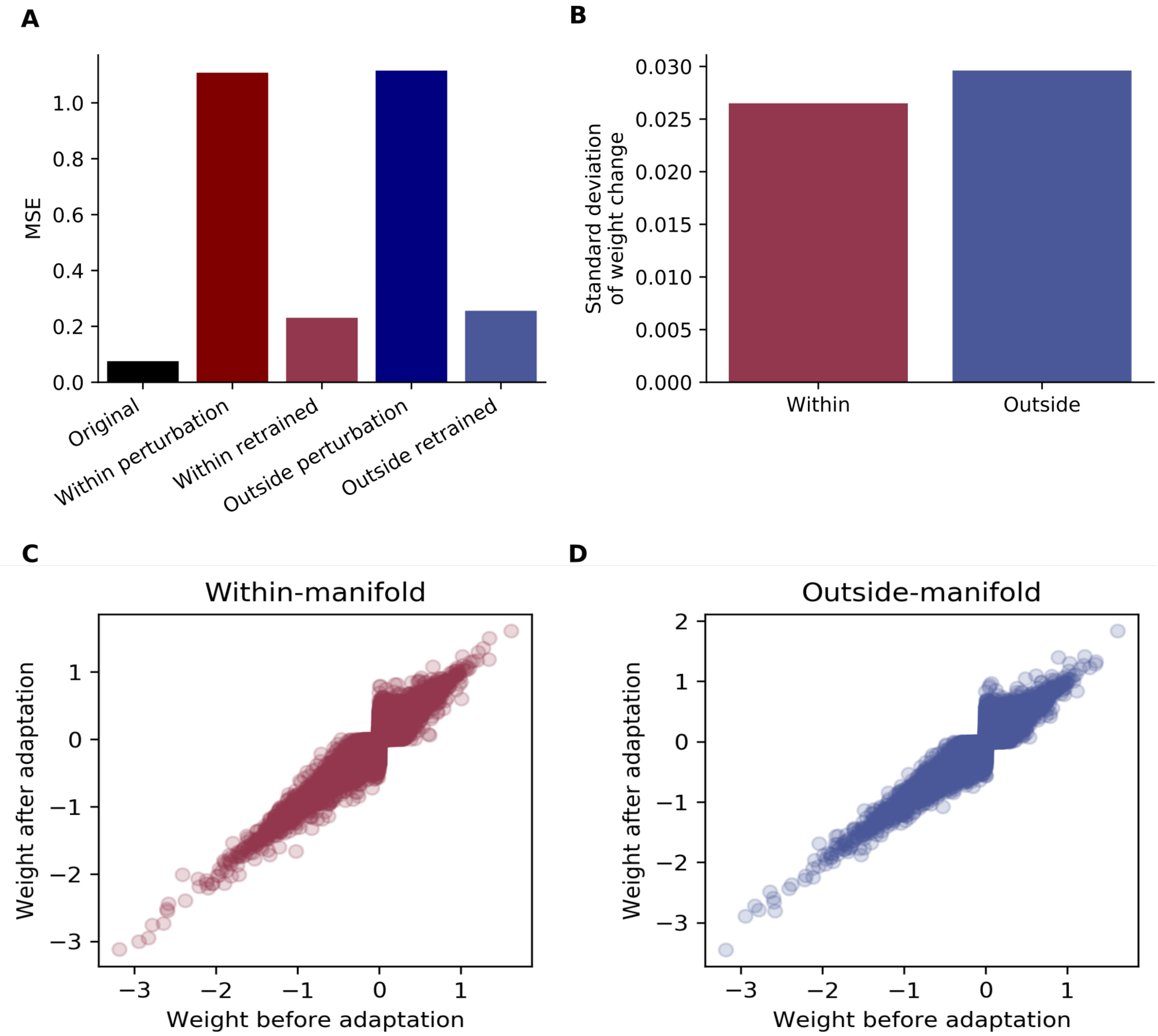


**Figure S5: Simulation with E-I network.** Simulation results for a network with Daleian connectivity. Each neuron has either positive (excitatory) or negative (inhibitory) outgoing weights. During learning, the sign of these weights is preserved. If a learning step would produce a sign flip we instead clip the weight to zero. (A) Training performance measured as mean squared error (MSE). (B) Standard deviation of weight change during within- and outside-manifold training. (C)&(D) Weights before versus after within- (C) or outside- (D) manifold training.

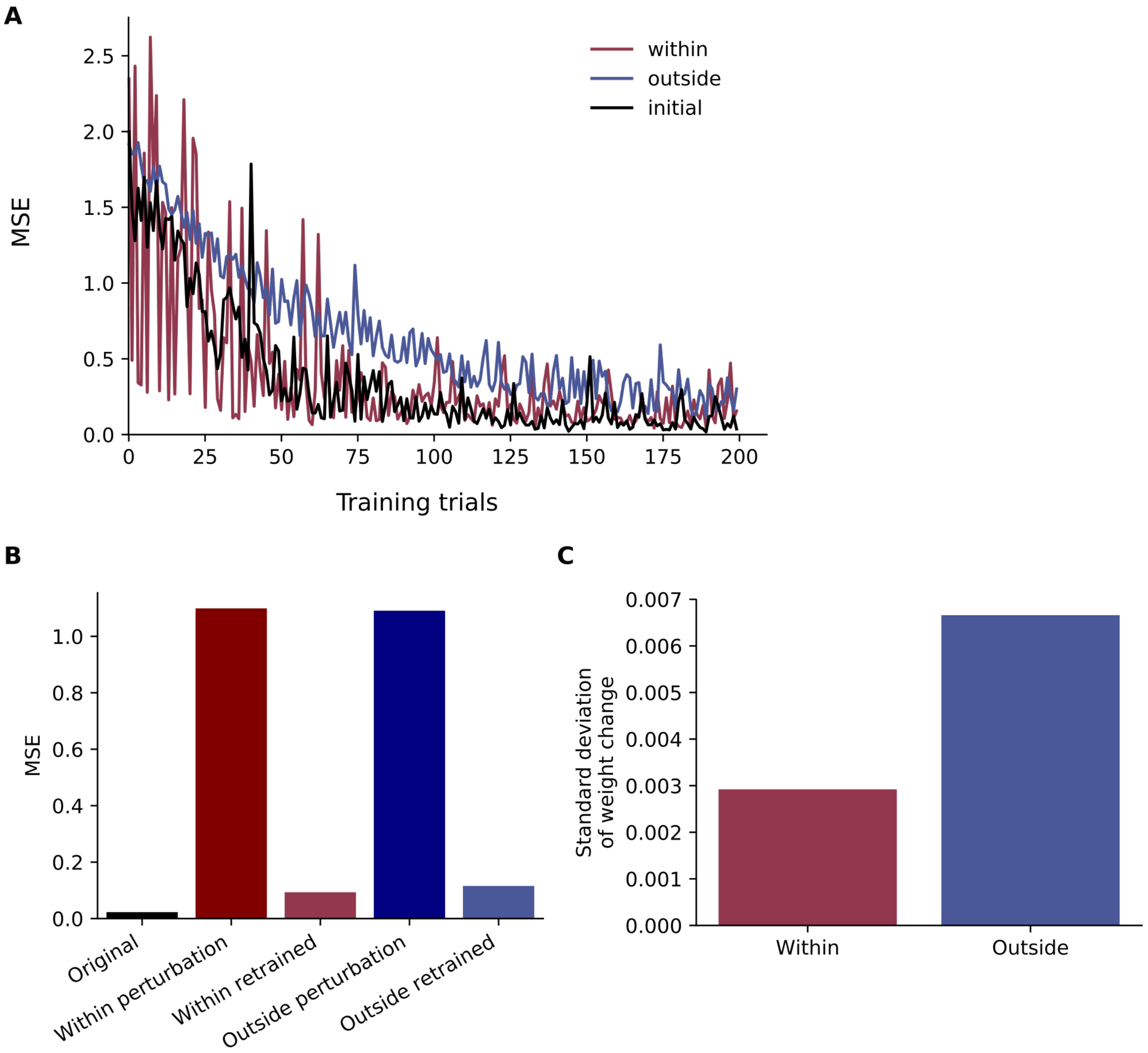


**Figure S6: Simulation with local learning rule.** Implementation of recently proposed local learning rule approximating backpropagation-through-time algorithm [Bellec et al., 2019]. The weight update is given by $dW_{ij}^t = -e_j^t(1-\tanh^2(x_j^t))\sum_{t'\leq t-1} r_i^{t'}$, where $e_j$ is the error for neuron $j$, $x_j$ is the activity of neuron $j$ and $r_i = \tanh(x_i)$ is the rate of neuron $i$. (A) Learning curves for initial, within- and outside-manifold training period. (B) Performance results after training measured as mean squared error (MSE). (C) Standard deviation of weight change during within- and outside-manifold training.

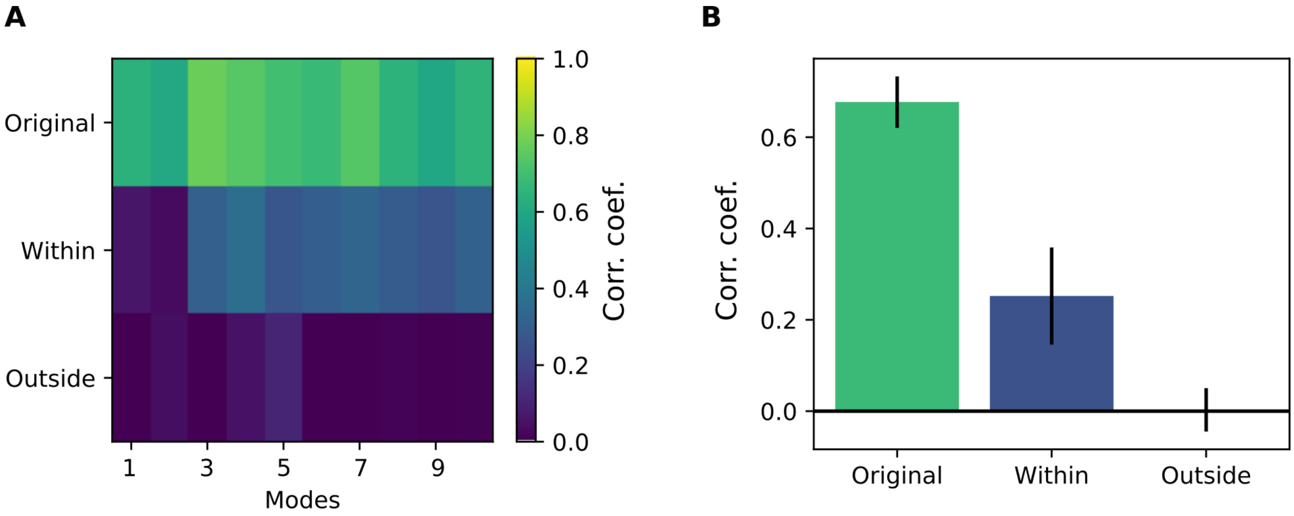


**Figure S7: Mode correlation between before and after retraining.** (A) Correlation between the neural modes before and after retraining for each of the six targets. Note, the neural manifold is in this case not the actual internal manifold (which is different after retraining), but the static one defined by the initial BCI mapping. We averaged over 20 simulations and the 6 targets. (B) Average mode correlation. Error bars are standard deviation across the ten modes.

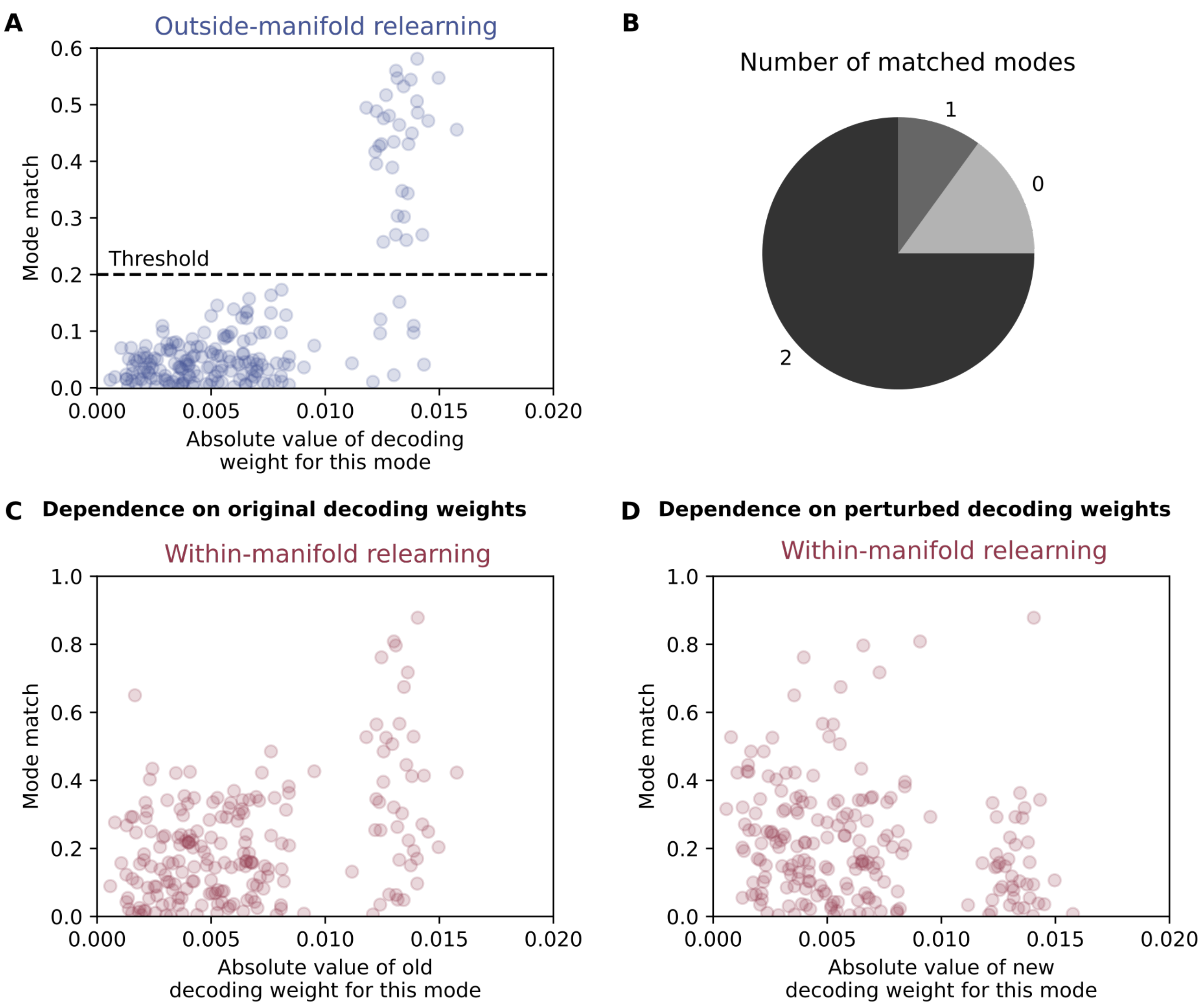


**Figure S8: Mode alignment during outside-manifold (A-B) and within-manifold (C-D) relearning.** (A) Mode match is defined as scalar product between the Eigenvector of the specific Eigenvalue before and after retraining. Each dot represents the mode match of one of ten Eigenvalues considered, and shows the result for one of 20 simulation runs (total 200 points shown). We defined a mode as matched if the scalar product is bigger than 0.2. (B) We quantified how many modes are matched per simulation run, taking the definition from (A). (C) Same as in (A), but now for within-manifold retraining. As the decoding weights are shuffled during a within-manifold perturbation one can compare the mode match to the decoding weights before and after perturbation. Here, we compared to before perturbation. (D) Mode match for within-manifold retraining compared to decoding weights after perturbation.

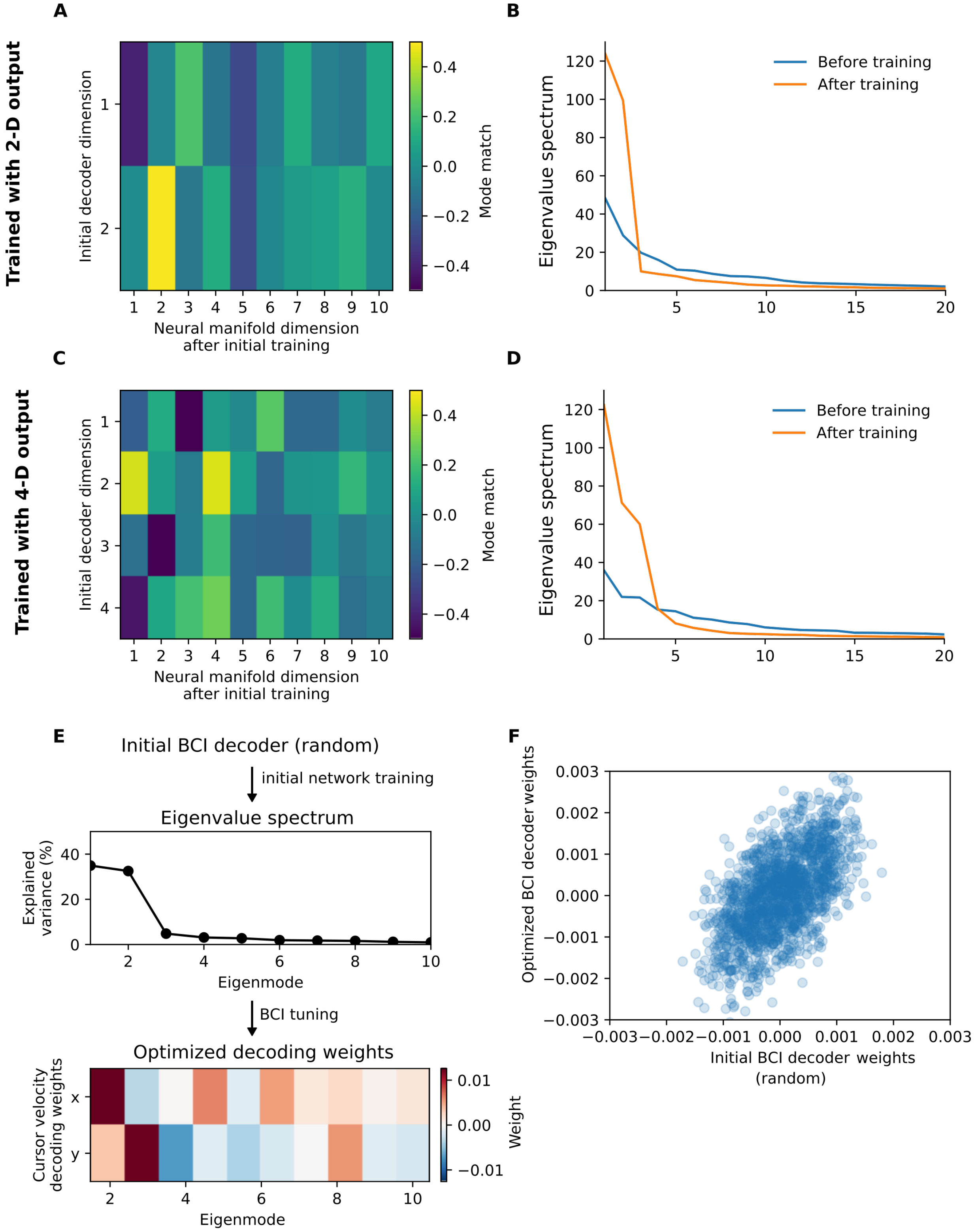


**Figure S9: Initial random decoder shapes neural manifold and final BCI mapping.** (A-B) Standard case where initial random decoder output is two-dimensional. (A) Mode match is calculated as scalar product between vectors. The initial decoder induces the first two strong neural modes, which are strongly amplified in the Eigenvalue spectrum after initial training (B). (C-D) Test case where initial random decoder output is four-dimensional. Here, the assignment of initial decoder and neural manifold modes is less prominent, meaning that there is no one mode specifically responsible for one output dimension. (C). In this case the number of amplified modes does not correspond to the dimensionality of the initial decoder (D) (E) Initial random decoder shapes Eigenvalue spectrum during initial network training phase, and Eigenvalue spectrum in turn shapes optimized BCI decoding weights $D$. (F) Relation between optimized BCI decoder and initial random one used for initial network training.

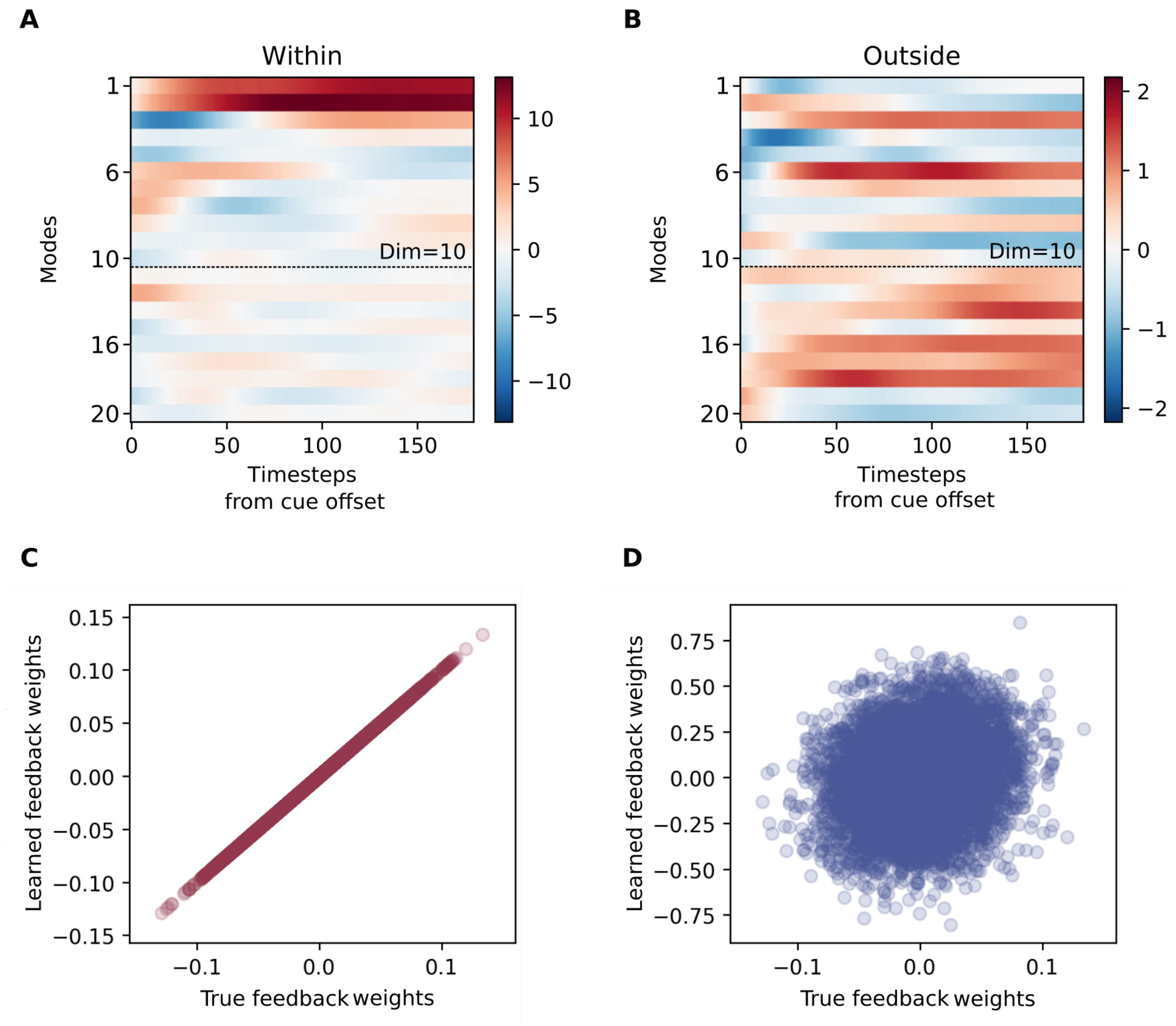


**Figure S10: Feedback learning on mode level.** (A-B) Mode activation during one example trial for a within-manifold perturbation (A) and an outside-manifold perturbation (B). Mode activation is obtained by projecting measured neural dynamics onto the original neural manifold, given by the transformation matrix $C$. (C-D) Results of linear regression to infer inverse of transformation matrix $C$ for a within-manifold perturbation (C) and an outside-manifold perturbation (D).

### Simplified model to investigate feedback learning

In order to understand why it is possible to learn feedback weights for a within-manifold perturbation, but not for an outside-manifold one, we developed and analysed a simplified setup. We created a synthetic dataset, which has no temporal structure, but is simply drawn from a multivariate Gaussian distribution. The number of variables was matched to the number of neurons in the simulated network, such that there was 800 random variables $\xi$ in total. Without loss of generality, we set the mean to zero, as we wanted to analyse the effect of the covariance structure on the ability to perform feedback learning. To create the covariance matrix $\mathbf{\Sigma}$ of the synthetic data, we firstly created the desired Eigenvalue spectrum which is given by

$$f(x;\gamma) = \exp^{-x/\gamma}$$

where $\gamma$ parametrizes how fast the spectrum decays and $x = 0$ correspond to the first Eigenvalue, $x = 1$ to the second and so forth. After setting the diagonal values of the covariance matrix, we performed a basis transformation which puts the synthetic data in the same Eigenspace as found in the simulated data.

$$\mathbf{\Sigma} = C^T \operatorname{diag}(f(0;\gamma), f(1;\gamma), ..., f(n-1;\gamma))C$$

$$\xi \sim \mathcal{N}((0, 0, ..., 0), \mathbf{\Sigma})$$

To simulate a within- or an outside-manifold BCI transformation, we used the Eigenvector space $C$ of the data and either took the first $d$ Eigenvectors (within-manifold) or the first $d$ Eigenvectors after shuffling the entries of each vector (outside-manifold). For Fig.S11 and S12 the number of modes $d$ - which are decoded - is set to 10 to match the number of modes used in the main paper.

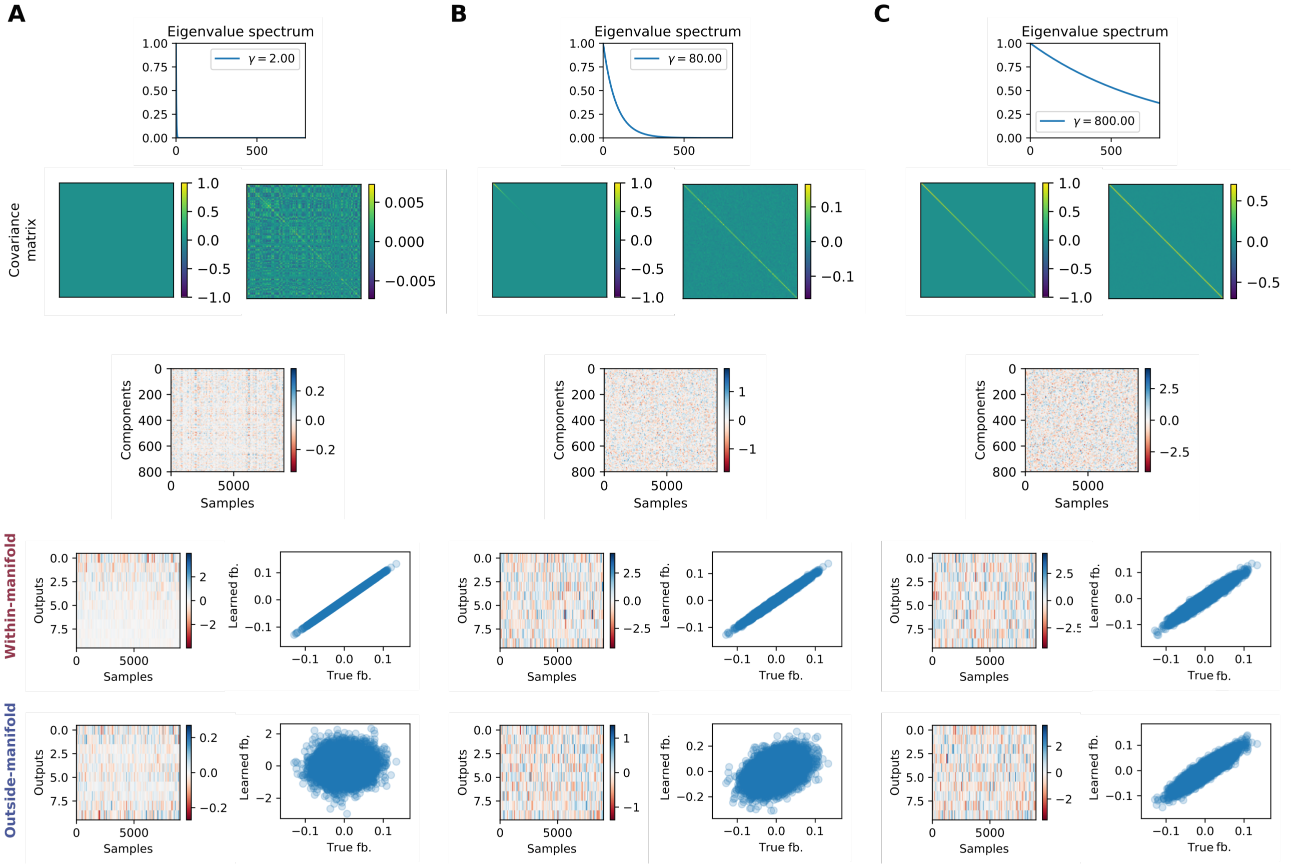


**Figure S11: Feedback learning on mode level for different Eigenvalue spectra.** (A) Narrow Eigenvalue spectrum. (B) Intermediate Eigenvalue spectrum. (C) Broad Eigenvalue spectrum. (Top panel) Imposed Eigenvalue spectrum. (Second panel from the top) Covariance matrix before and after basis transformation. (Third panel from the top) Synthesized random variables dataset. (Fourth and fifth panel from the top) Mode activation for a within-manifold (fourth) or outside-manifold (fifth) transformation and feedback weights learning result.

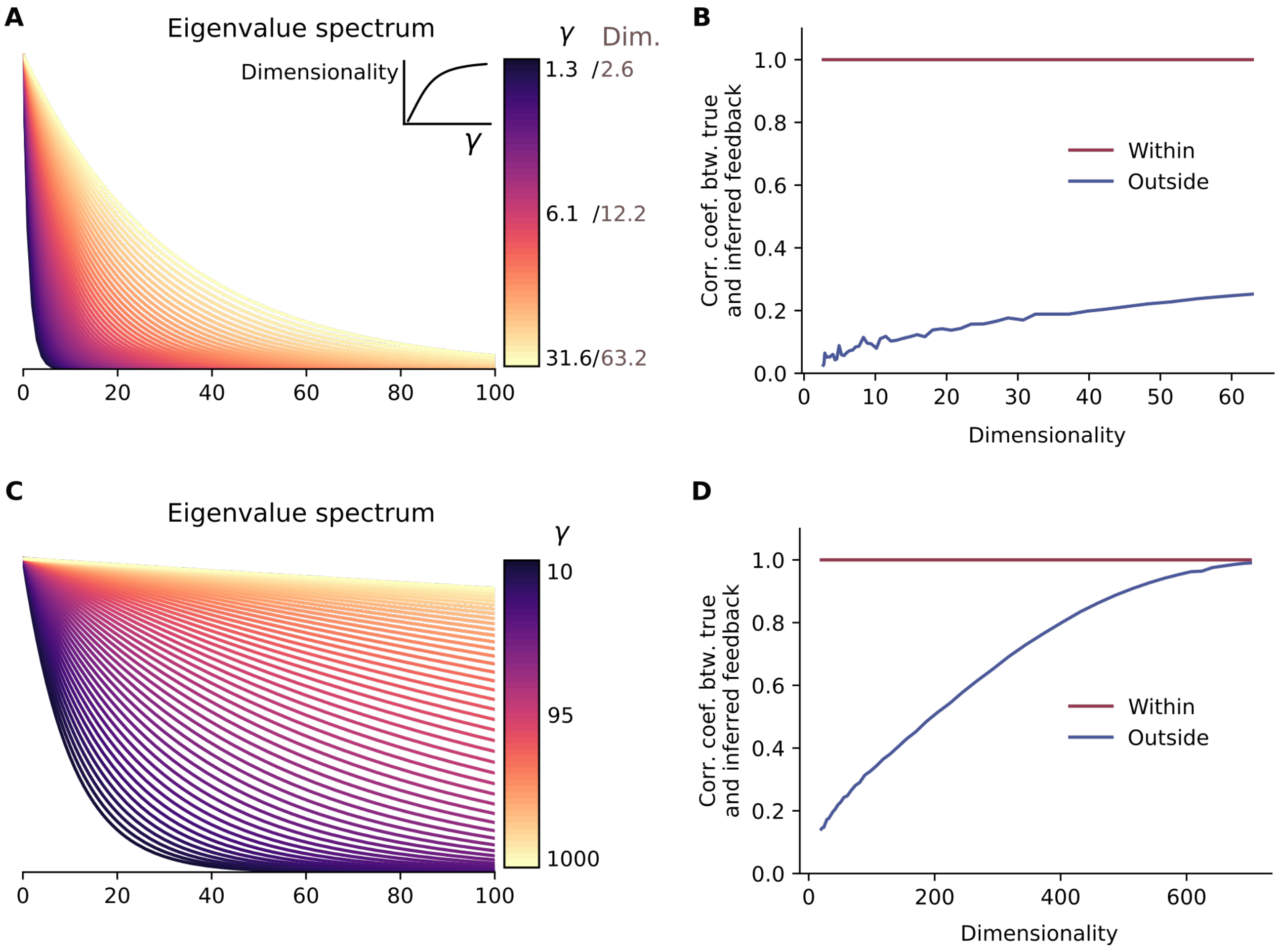


**Figure S12: Broader Eigenvalue spectrum improves feedback learning performance for outside-manifold perturbations.** (A)&(C) Imposed Eigenvalue spectra, defined by $f(x;\gamma) = \exp^{-x/\gamma}$. Dimensionality is calculated by $(\sum_i \lambda_i)^2 / \sum_i(\lambda_i^2)$ where $\lambda_i$ is the $i$th Eigenvalue. (B)&(D) Feedback learning results measured by correlation coefficient between inferred and true feedback weights, dependent on the dimensionality of the imposed Eigenvalue spectrum.
